# Regulation of itch-induced scratching by nucleus accumbens dopamine receptor expressing neurons

**DOI:** 10.64898/2026.04.16.718967

**Authors:** Jagat Narayan Prajapati, P B Nila, Debstuti Basu, Dipak K. Sahare, Dadasaheb M. Kokare, Arnab Barik

## Abstract

Scratching provides transient relief from itch, yet the neural circuit mechanisms that transform scratching into itch relief remain poorly understood. Midbrain dopaminergic neurons and their downstream targets in the lateral shell of the nucleus accumbens (NAc LaSh) are implicated in itch–scratch processing. Previous studies show that pharmacological manipulation of dopamine D1 and D2 receptors in the NAc LaSh alters scratching behavior, but the specific contributions of D1R- and D2R-expressing neurons during acute and chronic itch remain unclear. Here, we show that NAc LaSh^D1R^ ^and^ ^D2R^ neurons bidirectionally regulate scratching behavior across itch states. NAc LaSh^D1R^ neurons’ activity promotes scratching bouts, whereas NAc LaSh^D2R^ neurons preferentially facilitate scratch termination. Anterograde viral tracing revealed distinct brain-wide projection patterns of NAc LaSh^D1R^ ^and^ ^D2R^ neurons, which we functionally tested using projection-specific optogenetic manipulations. We found that NAc LaSh^D2R^ neurons terminate scratching by inhibiting neurons in the lateral parabrachial nucleus (LPBN), a key hub for itch processing. Furthermore, dopamine levels in the NAc LaSh were elevated during chronic itch compared with acute itch, suggesting enhanced dopaminergic signaling contributes to persistent scratching. Together, these findings identify circuit mechanisms linking reward pathways to itch regulation.

## Introduction

Scratching is an evolutionarily conserved behavioral response to itch sensation and serves as an important protective mechanism by suppressing itch through noxious mechanical stimulation and by promoting immune response against bacterial infections (Liu et al., 2025a). However, in cases of chronic itch, excessive scratching becomes maladaptive and significantly impacts quality of life (Carr et al., 2014; Sanders and Akiyama, 2018; Cevikbas and Lerner, 2020; Choragudi et al., 2023). Over the past few decades, considerable progress has been made in understanding the peripheral neural mechanisms underlying itch sensation (Dong and Dong, 2018; Lay and Dong, 2020; Guo et al., 2022; Ding et al., 2024; Prajapati et al., 2024). However, the central circuits underlying the itch-scratch cycle are relatively unexplored.

Prior human brain imaging studies using functional magnetic resonance imaging (fMRI) have shown that scratching-induced pleasantness activates brain reward system-associated areas, including the VTA and the nucleus accumbens (Papoiu et al., 2013; Mochizuki et al., 2014, 2015). However, these studies were limited by their temporal resolution, cell-type specificity, and mechanistic exploration. More recent studies in rodents have shown that VTA dopaminergic neurons are activated in a scratch-dependent manner, and inhibition of these neurons attenuates scratching-induced reward (Yuan et al., 2018; Su et al., 2019; Setsu et al., 2021). Activity of dopaminergic neurons in the VTA correlated with scratching and mediated reward associated with itch relief. Furthermore, it was found that scratch-induced activation of VTA dopaminergic neurons drives DA release in the NAc LaSh, but not in the NAc MeSh (Liang et al., 2022). NAc LaSh is an anatomically distinct but understudied subdivision of NAc, and is activated by reward and reward-predictive cues (Yang et al., 2018; de Jong et al., 2019; Chen et al., 2023). Interestingly, recent studies in mice suggest that pharmacological blockade of D1R signaling in NAc LaSh and activation of D2R signaling impair pruritogen-induced scratching behaviors, indicating an opposing role of D1R and D2R signaling in NAc LaSh in itch processing (Liang et al., 2022). These findings are in line with previous studies in rodents and primates, which showed that pharmacological manipulation of D1R and D2R receptors attenuated scratching (Merali and Piggins, 1990; Rosenzweig-Lipson et al., 1994; Pellón et al., 1995; Akimoto and Furuse, 2011). Nevertheless, the function of the neurons that express D1R and D2R receptors in the NAc LaSh (NAc LaSh^D1R^ and NAc LaSh^D2R^) during acute and chronic itch-induced scratching remains relatively unexplored.

Here, we used promoter-driven viral vectors to target D1R- and D2R-expressing neurons in NAc LaSh. We found that the NAc LaSh^D1R^ and NAc LaSh^D2R^ neurons were differentially engaged during the start and end of scratching bouts. We tested their activity during chloroquine-mediated acute and chronic psoriatic itch conditions and found that these neurons process acute and chronic itch in a similar manner. Observations from experiments measuring in vivo dopamine release dynamics indicated that the initiation and termination of scratching correlate with dopamine release onto NAc LaSh^D1R^ neurons, suggesting that alterations in dopamine levels may impair normal itch-scratching coupling. Indeed, we found that the dopamine levels are elevated in mice with psoriasis, and L-DOPA administration perturbed scratching responses to acute but not chronic pruritic stimuli. Further, we mapped the downstream targets of the NAc LaSh^D1R^ and NAc LaSh^D2R^ neurons and found that the brainstem lateral parabrachial nuclei (LPBN) are preferentially targeted by the NAc LaSh^D2R^ neurons. In agreement, stimulation of D2R terminals in the LPBN suppressed itch. Thus, in this manuscript, we have furthered our understanding of how midbrain dopaminergic neurons play a critical role in relieving itch and how this feedback mechanism goes awry in pathological conditions.

## Methods

### Mouse line

Animal care and experimental procedures were performed in accordance with protocols approved by the CPSCEA at the Indian Institute of Science. CD-1 mice (both males and females, 7-12 weeks old) were purchased from the Central Animal Facility (CAF) at IISc. The animals were housed at the CAF under standard animal housing conditions, with a 12-hour light-dark cycle and *ad libitum* access to food and water. An equal number of males and females were used in all the behavioral experiments. All mice used in the behavioral assays were between 7 and 12 weeks old. All behaviors occurred during the light cycle.

A microdialysis study to estimate dopamine concentration was done on male Wistar rats. Adult male Wistar rats (250-300 g) were group-housed under controlled conditions and 12:12 h light/dark cycle (lights on 07:00 AM). Food pellets (Amrut Laboratory Animal Feeds, Pune, India) and water were provided *ad libitum*. Behavioral tasks were carried out from 8:00 to 12:00 under strict compliance of the Institutional Animal Ethics Committee (IAEC/UDPS/2025/01/01) constituted at the Department of Pharmaceutical Sciences, RTM Nagpur University, India. Study complied with the guidelines of the Committee for Control and Supervision of Experiments on Animals (CCSEA), Ministry of Fisheries, Animal Husbandry and Dairying, Government of India (Registration no. 92/PO/Re/S/1999/CCSEA).

### Viral Vectors

Vector used and sources: pAAV5-FLEX-tdTomato (Addgene, Catalog# 28306-AAV1, titer- 1.6 x 10^13^ GC/ml), pAAV5-hsyn-DIO-hM3D(Gq)-mCherry (Addgene, Catalog# 44361, titer-1.8 x 10^13^ GC/ml), AAV9.syn.flex.GcaMP8s (Addgene, Catalog# 162377, titer- 2.7 x 10^13^ GC/ml), rAAV2/9-hSyn-DIO-hM4DIGi)-eYFP (BrainVTA, Catalog# PT-0344, titer- 5.34 x 10^12^ vg/ml), rAAV2/9-D1R-CRE-WPRE-hGH-polyA (BrainVTA, Catalog# PT-1217, titer- 4.72 x 10^12^ vg/ml), rAAV2/9-D2R (ENK)-CRE-WPRE-bGH-polyA (BrainVTA, Catalog# PT-0571, titer- 2 x 10^12^ vg/ml), rAAV2/5-EF1α-DIO- dLight1.2 (BrainVTA, Catalog# PT-1296, titer- 1.18 x 10^13^ vg/ml), AAV5-DIO-ChR2-mCherry (Addgene, Catalog# 20297, titer- 1.4 x 10^13^ GC/ml), AAV9/2-hSyn-DIO-mSyp1_EGFP(University of Zurich, Catalog# v484-9, titer- 7.8 x 10^12^ vg/ml).

### Antibodies

Chicken anti-GFP antibody (aveslabs catalog# 1010), Phospho-c-Fos (Ser32) Rabbit monoclonal antibody (Cell Signaling Technology Catalog# 5348), Goat anti-tdTomato (SICGEN catalog# AB8181), Goat anti-Chicken IgY (H+L) secondary Antibody, Alexa Fluor™ 488 (Invitrogen catalog# A11039), Donkey anti-Rabbit IgG (H+L) secondary Antibody, Alexa Fluor™ 488 (Invitrogen catalog# A21206), Donkey anti-Goat IgG (H+L) secondary Antibody, Alexa Fluor™ 594 (Invitrogen catalog# A11058),

### Stereotaxic injections

Mice were anesthetized with 2% isoflurane/oxygen before and during the surgery and mounted on the stereotaxic frame (RWD 69100 Rotational Digital Stereotaxic Frame). An incision was made to expose the skull, and subsequently, the skull was aligned to the horizontal plane. A craniotomy was performed at the marked point using a handheld microdrill (RWD). A Hamilton syringe (10 μL) with a glass-pulled needle was used to infuse 300 nL of viral particles at a rate of 100 nL/min. Viral particles were prepared by mixing 1 µL of either D1R-Cre or D2R-Cre expressing AAVs with 1 µL of Cre-dependent AAVs expressing any of these transgenes: hM3Dq, hM4Di, GCaMP8s, dLight1.2, and ChR2 to drive the expression of these transgenes in either D1R neurons or D2R neurons. This 1:1 mixture of viral particles was further diluted with 1 µL of saline, and the resulting viral particles were infused (300 nL at a rate of 100 nL/min) into the nucleus accumbens lateral shell (NAc LaSh) at the following coordinates: AP: 1.02; ML: ±1.75; DV: −4.60.

### Microdialysis for estimation of dopamine (DA) and 3,4-Dihydroxyphenylacetic acid (DOPAC) in the lateral shell of the nucleus accumbens

The concentrations of DA and its metabolite (DOPAC) were assayed in the dialysate collected from the different groups of rats using a microdialysis sampling technique and quantified using a high-performance liquid chromatography-electrochemical detector (HPLC-ECD). The method has been standardized in our laboratory (Awathale et al., 2021; Kawade et al., 2022). Using the stereotaxic procedure described by Awathale et al. (2021), the rats were unilaterally implanted with a vertical guide cannula (BASi Instruments, USA, Cat. no. MD-2251) targeted at the lateral nucleus accumbens shell (LAcbSh), the (+1.0 mm anterior, -2.9 mm mediolateral and -6.5 mm ventral with reference to bregma from skull surface). Following a 7-day recovery period, the rats were subjected to imiquimod (5% w/w, 0.25 g/day) treatment for five days, and thereafter, on day 12, the dummy cannula targeted at the LAcbSh was replaced with the microdialysis probe (2 mm membrane length, OD 320 μm and ID 220 μm, BASi systems, USA, Cat. no. MD-2200) pre-saturated with aCSF. The microdialysis probe extended 2 mm beyond the tip of the guide cannula up to 8.5 mm. The timeline of the surgical procedure, imiquimod treatment and dialysate collection are summarized in Fig. 4A. The probe was perfused with aCSF (140 mM NaCl, 3 mM KCl, 2.4 mM CaCl_2_, 1 mM MgCl_2_, 1.2 mM Na_2_HPO_4_, 0.27 mM NaH_2_PO_4_, 7.2 mM glucose, pH 7.4, all reagents Merck Life Science Pvt. Ltd., India) using gastight syringe (MDN-0100, BASi systems, USA) operated by infusion pump (MD-1001, BASi system, USA) at a flow rate 1 μl/min. The microdialysis probe was inserted in the guide cannula, and two hours were allowed for stabilization. Thereafter, dialysates were collected for 20 min intervals (Fig. 4A). The dialysate collection vials were prefilled with 10 μl of perchloric acid (0.2 M, Sigma, USA, Cat. no. 50439) and mounted in a refrigerated fraction collector (MD-1201, BASi systems, USA, temperature 0 °C).

The collected samples were stored at 0 °C, and the concentrations of DA and DOPAC in the dialysates were analyzed using a HPLC-ECD system (Waters Pvt. Ltd., USA, Model no. 2456). The Ascentis C18 column (25 cm × 4.6 mm, 5 µm) (Sigma Aldrich-Supelco, Cat. no. 581325-U) was used. Mobile phase consisted of 75 mM sodium phosphate monobasic, 350 mg/litter 1-octanesulfonic acid sodium salt, 0.5 mM EDTA, 1% tetrahydrofuran and 15% acetonitrile at pH 3.0 (adjusted with 85% phosphoric acid) (all reagents Sigma, USA), while flow rate (1 ml/min), oxidizing potential (+0.70 V), and run-time was maintained at 10 min (Li et al., 2002; Deehan et al., 2013). The retention time (RT) of dialysate samples was compared with that of the external standards of DA (RT: 3.204 min, 5 pg/µl) and DOPAC (RT: 5.041 min, 15 pg/µl) (Fig. 4C). The biological samples were injected, and the area under the curve (AUC) was recorded. The AUC was automatically calculated (Breeze^TM^2 software, Waters Pvt. Ltd., USA). After each experiment, the functionality of the microdialysis system was assessed with a K^+^ stimulation (60 mM) experiment. In brief, three baseline samples were collected by infusing aCSF through the probe. Thereafter, 60 mM K^+^ (equimolar to Na+) was applied via probe for 30 min, and corresponding samples were collected at 10 min intervals at each time point (Kawade et al., 2022). Only the rats with the correct probe location were included in the statistical test.

### Fiber optic cannula implantation

A fiber optic cannula from RWD (Ø1.25 mm Ceramic Ferrule, 300 μm Core, 0.39NA, L = 6 mm, catalog# R-FOC-BL300C-39NA) was implanted at AP: 1.02, ML: ±1.75; DV −4.60 in the NAc LaSh of the GCaMP8s or dLight1.2 infused mice. The cannula was fixed to the skull using light-cured dental cement (GC corporation powder- catalog# 002505, liquid- catalog# 002524). Animals were allowed to recover for at least 1 week before performing behavioral tests.

Successful labeling and fiber implantation were confirmed post hoc by GFP staining, which indicated viral expression and fiber-induced injury, respectively. Only animals with viral-mediated gene expression and fiber implantations at the intended locations, as observed in post hoc tests, were included in the analysis.

### Fiber photometry

A dual-channel fiber photometry system from RWD (R810) was used to collect the data(Gunaydin et al., 2014; Kim et al., 2016). The light from two light LEDs (410 and 470 nm) was passed through a fiber optic cable (RWD- Ø1.25 mm Ceramic Ferrule, 200 μm Core, 0.39NA, L = 2 m, catalog# R-FC-L-N3-200-L1) coupled to the cannula implanted in the mouse. Fluorescence emission was acquired through the same fiber-optic cable and directed onto a CMOS camera via a dichroic filter. Mice were lightly anesthetized, and the fiber-optic cable was connected to the optical cannula attached to the mouse skull. Mice were habituated to the fibers for 2 days before performing any behavioral assays. The output power was adjusted to 30%, resulting in a power range of 20-50 µW at the fiber tip. The signals were acquired at 100 frames per second. The data was analyzed using the RWD photometry software, and .csv files were generated. The start and end of scratching were timestamped. Due to variability in the scratching bout duration, we plotted the 3 seconds before and after the scratch start and scratch end separately. All trace graphs were plotted from .csv files using GraphPad Prism software version 8.

### Chemogenetic activation and inhibition

For chemogenetic activation or inhibition of NAc LaSh D1R or D2R neurons, deschloroclozapine (DCZ) (Hello Bio, catalog# HB9126), at a dose of 2 µg/kg body weight, was administered intraperitoneally (i.p.) into the mice(Nagai et al., 2020). All behavioral assays were performed 15 minutes after DCZ administration.

### L-DOPA administration

L-DOPA (Sigma, catalog# D9628) was freshly prepared for each experiment by dissolving it in saline. Mice were i.p. administered with L-DOPA at doses of 5 or 10 mg/kg body weight prior to the behavioral experiment. Behavioral assays were performed at two time points: 15 min and 2.5 h after L-DOPA administration.

### Optogenetic activation

For optogenetic activation of NAc LaSh^D1R^ or NAc LaSh^D2R^ neurons using channelrhodopsin bilateral fiber optic cannula (Ø1.25 mm Ceramic Ferrule, 300 μm Core, 0.39NA, L = 6 mm, catalog# R-FOC-BL300C-39NA) was implanted at NAc LaSh (AP: 1.02, ML: ±1.75, DV: -4.60). One week after the implantation, mice were habituated in a fresh cage with a fiber optic cable (RWD- Ø1.25 mm Ceramic Ferrule, 200 μm Core, 0.39NA, L = 2 m, catalog# R-FC-L-N3-200-L1) for 2 days and then used for behavioural experiments. The IOS-465 Intelligent Optogenetics System (RWD) was used to deliver 465nm laser pulses (20Hz, 10 mW, duration: 5ms, interpulse duration: 10ms). Fiber optic cannula-implanted mice were briefly anesthetized using isoflurane, and an optical fiber was connected to the fiber optic cannula to deliver laser light. The scratching behavior of mice was recorded with a 5-minute light-on and a 5-minute light-off cycle for 1 hour.

### Behavioural assays

#### Chloroquine-induced itch assay

The nape of the neck of mice was shaved with a hand-held Philips shaver 2-3 days before behavioral experimentation, and the mice were habituated in the behavior room. Unless otherwise stated, the mice used for behavioral studies were blinded prior to initiation of the studies by an individual not involved in the experimentations described here. All behavioral experiments were quantified by one experimenter and randomly cross-verified by another. All itch experiments were videotaped with a Logitech camera, and videos were acquired through vendor-supplied software. Mice were individually placed in four-part plexiglass chambers with dimensions of 6 cm × 6 cm × 14 cm. The roof of the chamber had holes for air ventilation.

Animals were habituated in the chamber for 15 min before chloroquine injections. DCZ 2 µg/kg body weight was administered intraperitoneally (i.p.) 15 min before chloroquine injection. Chloroquine (375 µg/75 µl) (Sigma Catalog# C6628) was administered intradermally into the nape of the neck of the mice, and the subsequent scratching behavior was recorded for 30 min (Prajapati et al., 2025). Hind leg-directed scratching of the nape was characterized as a scratch, and the videos were quantified, blinded to the experimental conditions.

### Imiquimod induced psoriatic itch

Imiquimod (5% w/w from Glenmark) was used to induce psoriasis in mice (van der Fits et al., 2009; Sakai et al., 2016). To induce psoriasis-like chronic itch, the nape of each mouse was shaved using Veet hair removal cream, and imiquimod was topically applied once daily to the shaved area for six consecutive days. This treatment reliably induced inflamed, scaly skin lesions characterized by thickened and dry epidermis. Mice were manually inspected, and only those exhibiting pronounced psoriatic features- namely, inflamed, thickened, and scaly skin-were selected for behavioral analysis. Following psoriasis induction, mice were individually placed in four-compartment plexiglass chambers for habituation, after which their behavior was recorded for 30 minutes. Spontaneous scratching was defined as hind limb-directed contact to the nape region. Scratching bouts were quantified by observers blinded to the experimental conditions.

### Immunostaining, multiplex in situ hybridization, and confocal microscopy

Mice were anesthetized with isoflurane and perfused transcardially with 1X Phosphate Buffered Saline (PBS) (Takara catalog# T9181) and 4 % Paraformaldehyde (PFA) (Ted Pella, Inc. catalog# 18505). Harvested brains were further fixed in 4% PFA overnight, then transferred to 15% and 30% sucrose for serial dehydration. Brain tissues were placed in the Cryo-Embedding Compound (Ted Pella, Inc.) and frozen at -40°C. Subsequently, 50 µm-thick coronal brain sections were cut using a cryostat (RWD Minux FS800). For immunostaining experiments, tissue sections were rinsed in 1X PBS (3 times) and incubated in the blocking buffer (5 % Bovine Serum Albumin (BSA) + 0.5 % Triton X-100 + 1X PBS) (BSA- HIMEDIA catalog# TC194, Triton X-100 SRL catalog# 64518) for one hr at room temperature. Sections were then incubated in the primary antibody (dilution 1:1000 X in blocking buffer) at room temperature overnight. Sections were rinsed 3 times with 1X PBS + 0.5 % Triton X-100 solution and incubated for two hr in Alexa Fluor conjugated goat anti-rabbit/ chicken or donkey anti-goat/rabbit secondary antibodies (dilution 1:1000 X in blocking buffer) along with DAPI (SRL catalog# 18668) at room temperature. Then, sections were washed with 1X PBS + 0.5% Triton X-100 and mounted onto charged glass slides (Ted Pella, Inc., catalog #260382-3). Citifluor AF-1 mounting media (Ted Pella, Inc. catalog# 19470-1) was used to cover-slip (Blue star microscopic cover glass 24 x 60 mm 10 Gms) the slides. Subsequently, sections were imaged on the upright fluorescence microscope (Khush Enterprises, Bengaluru) (2X, 4X, and 10X lenses) and a Confocal Microscope (Leica SP8 Falcon, Germany). ImageJ/FIJI processing software was used to process the images. Confocal images were processed using the Leica Image Analysis Suite.

For *in situ* hybridization (ISH), fresh brains were rapidly harvested and flash-frozen at – 80C°C. Coronal sections (20 µm) were prepared using a cryostat. Multiplex ISH was performed using the manual RNAscope assay (Advanced Cell Diagnostics, ACD). Target-specific probes were obtained from the ACD online catalog: *drd1* (Ref. #461901), *tdTomato* (Ref. #317041), and *drd2* (Ref. #406501). Frozen brain sections were fixed in 4% paraformaldehyde (PFA; Ted Pella, Inc., Cat. #18505) for 15 minutes at room temperature, followed by sequential dehydration in graded ethanol (Hayman, Cat. #64-17-5) for 20 minutes. After brief air drying, a hydrophobic barrier was drawn around each tissue section. RNAscope Hydrogen Peroxide (Cat. #322335) was applied for 10 minutes, followed by two rinses in nuclease-free water (MP Biomedicals, Cat. #112450204). Sections were then incubated with Protease IV (Cat. #322336) for 30 minutes, followed by rinsing twice in nuclease-free water. A mixture of probes was prepared at a 50:1 ratio for *drd1* or *drd2* (channel 1) and *tdTomato* (channel 2), and applied to the sections. Hybridization was performed for 2.5 hours at 40°C using the HyperChrome hybridization system (HyperChrome, Cat. #EHP 500AS). Following hybridization, sections were washed with 1X wash buffer (Cat. #320058), and signal amplification and chromogenic development were performed according to the manufacturer’s protocol (ACDBIO). Images for anatomical analysis were acquired using 10X and 20X objectives on the upright fluorescence microscope (Khush Enterprises, Bengaluru).

### Brainwide projections mapping of NAc LaSh D1R or D2R neurons

To map the brain-wide projections of the NAc LaSh^D1R^ or NAc LaSh^D2R^ neurons, AAV-DIO-mSynGFP, along with either D1R- or D2R-Cre expressing AAV, was injected into the NAc of CD1 mice. 3 weeks later, brain tissues were harvested, and the brain-wide projections of the NAc LaSh^D1R^ or NAc LaSh^D2R^ neurons were imaged under fluorescent microscopy. Gain and exposure time were kept constant throughout the imaging session. A box area was selected within the region containing the SynGFP puncta to calculate the projection density, and the mean intensity (F_Total_) was measured in ImageJ/FIJI. To calculate the background intensity (F_Background_), the same box area was dragged to a region devoid of SynGFP puncta on the same brain section, and the mean intensity was then calculated. The mean fluorescent intensity of the region of interest (F_ROI_) was reported as F_ROI_= F_Total_ - F_Background_.

### Quantification and statistical analysis

All statistical analyses were performed using GraphPad PRISM 8.0.2 software. Student t-test and Two-way ANOVA tests were performed wherever applicable. ns > 0.05, ∗ P ≤ 0.05, ∗∗ P ≤ 0.01, ∗∗∗ P ≤ 0.001, ∗∗∗∗ P ≤ 0.0005.

## Results

### Engagement patterns of NAc LaSh^D1R^ and NAc LaSh^D2R^ neurons during acute and chronic psoriatic itch

Recent studies indicate that D1R and D2R receptors in the NAc LaSh and the dopamine signaling in these neurons that express them may play a crucial role in acute itch processing (Liang et al., 2022; Inokuchi-Sakata et al., 2024). However, it remains unclear how the activity of NAc LaSh^D1R^ and NAc LaSh^D2R^ neurons correlates with scratching evoked by acute itch and whether this correlation evolves with pathological itch. To address this gap, we deployed adeno-associated viral (AAV) vectors expressing Cre recombinase under the control of the D1R or D2R promoter, thereby specifically targeting D1R- and D2R-expressing neurons (Liu et al., 2025b; Zhang et al., 2025). This approach enables selective genetic targeting of D1R or D2R neurons in the NAc LaSh (Fig. S1). First, we validated the efficiency of these viral vector constructs for labeling distinct neuronal populations. To that end, we co-injected AAV-D1R-Cre with a Cre-dependent tdTomato expressing viral vector (AAV-DIO-tdTomato) into the NAc LaSh of wild-type CD1 mice (Fig. S1A). We performed multiplexed *in sit*u hybridization (RNAscope) on the NAc targeting the *Drd1* and *tdTomato* genes, and quantified the overlap between the *Drd1* and *tdTomato. We found* that the AAV-D1R-Cre viral vector labeled approximately 88% of the D1R-expressing neurons (Fig. S1B and C), suggesting that AAV-D1R-Cre effectively targeted the D1R neuronal population in the NAc LaSh. Similarly, we tested the efficiency of the AAV-D2R-Cre viral vector and found a labeling efficiency of 74.57% (Fig. S1D-F).

Next, to investigate neural activity in NAc LaSh^D1R^ neurons during acute itch-induced scratching, we expressed GCaMP8s (Zhang et al., 2023), a genetically encoded calcium sensor, selectively in NAc LaSh^D1R^ neurons using the same AAV viral vector (Fig. S1). Two viral vectors, AAV-DIO-GCaMP8s and AAV-D1R-Cre, were co-infused with the help of a mouse stereotactic device coupled with a microinfusion pump into the NAc LaSh of wild-type CD1 mice (Fig. 1A). After two weeks, a fiber optic cannula was implanted just above the NAc LaSh to record GCaMP8s fluorescence via fiber photometry (Gunaydin et al., 2014; Kim et al., 2016).

**Fig. 1.**
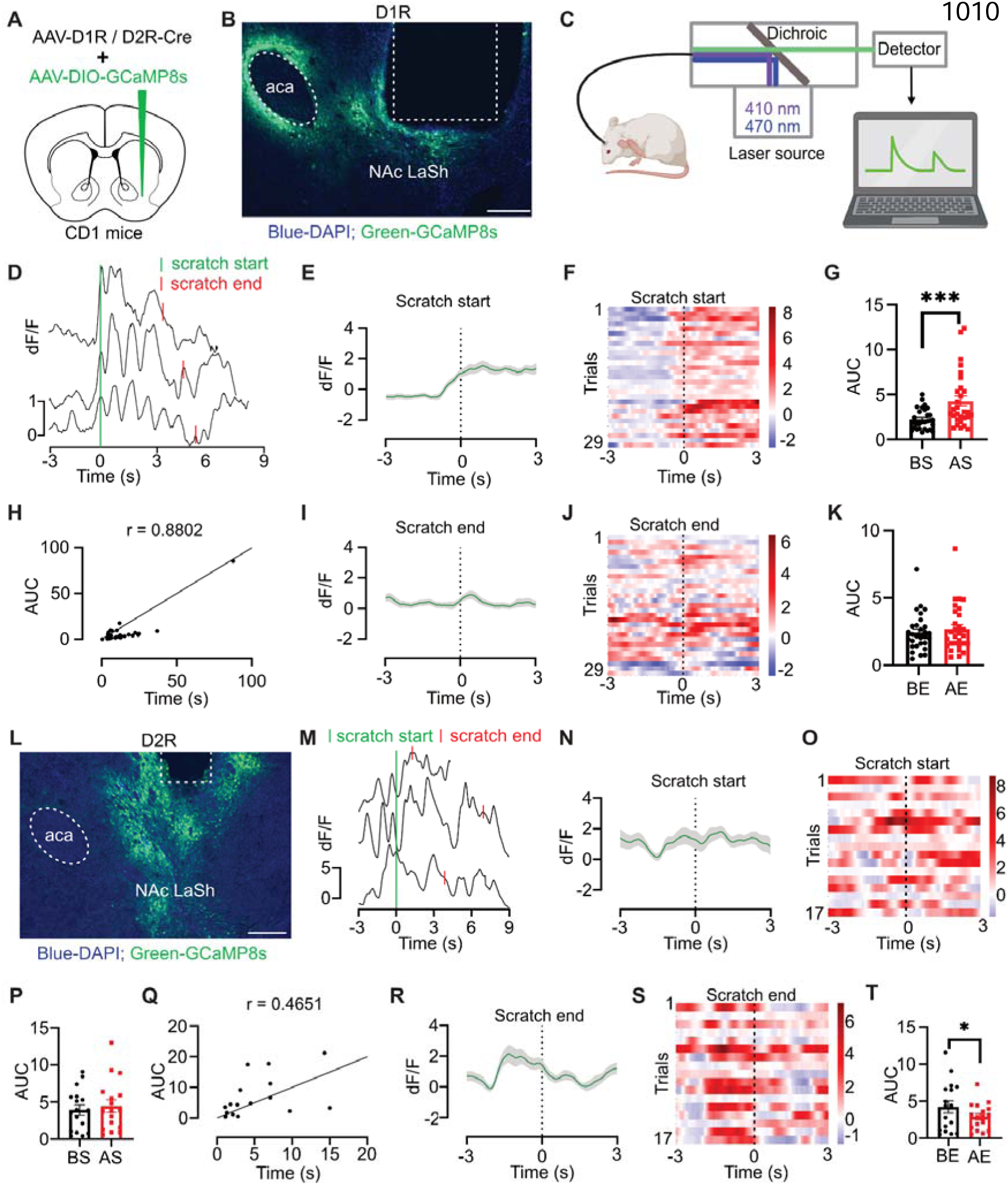
Engagement patterns of NAc LaSh^D1R^ and NAc LaSh^D2R^ neurons during acute itch. (A) Schematic of virus injection in the NAc LaSh. AAV-D1R-Cre or AAV-D2R-Cre, and AAV encoding Cre-dependent GCaMP8s were injected into the NAc LaSh of CD1 mice. (B) Coronal section image of the NAc LaSh showing D1R-dependent expression of GCaMP8s (green; DAPI-blue) and placement of the optic fiber cannula, scale bar 200 µm. (C) Schematic showing calcium activity recording using a fiber photometry system while the mouse is engaged in scratching behavior. (D) Representative ΔF/F traces of NAc LaSh^D1R^ neurons from three trials aligned to the start of chloroquine-evoked scratching. (E) The average fluorescence from NAc LaSh^D1R^ neurons at the start of scratch, with shaded area indicating SEM (n = 29 trials from 2 mice). (F) Heatmap of the fluorescence signal from NAc LaSh^D1R^ neurons at the start of scratch. (G) Quantification of area under the curve (AUC) of NAc LaSh^D1R^ activity 3 seconds before (BS) and after (AS) the start of scratch (t-test, 2.039 ± 0.4827 ***P = 0.0002, n = 29 trials). (H) Correlation of AUC of NAc LaSh^D1R^ activity with scratching duration (****P < 0.0001). (I) The average fluorescence from NAc LaSh^D1R^ neurons at the end of scratch, with shaded area indicating SEM (n = 29 trials from 2 mice ). (J) Heatmap of the fluorescence signal from NAc LaSh^D1R^ neurons at the end of scratch. (K) Quantification of area under the curve (AUC) of NAc LaSh^D1R^ activity 3 seconds before (BE) and after (AE) the end of scratch. (L) Coronal section image of the NAc LaSh confirming the D2R-dependent expression of GCaMP8s (green; DAPI-blue) and placement of the optic fiber, scale bar 200 µm. (M) Representative ΔF/F traces of NAc LaSh^D2R^ neurons from three trials aligned to the start of chloroquine-evoked scratching. (N) The average fluorescence from NAc LaSh^D2R^ neurons at the start of scratch, with shaded area indicating SEM (n = 17 trials from 2 mice). (O) Heatmap of the fluorescence signal from NAc LaSh^D2R^ neurons at the start of scratch. (P) Quantification of area under the curve (AUC) of NAc LaSh^D2R^ calcium activity 3 seconds before (BS) and after (AS) the start of scratch. (Q) Correlation of AUC of NAc LaSh^D2R^ activity with scratching duration. (R) The average fluorescence from NAc LaSh^D2R^ neurons at the end of scratch, with the shaded area indicating SEM (n = 17 trials from 2 mice). (S) Heatmap of the fluorescence signal from NAc LaSh^D2R^ neurons at the end of scratch. (T) Quantification of area under the curve (AUC) of NAc LaSh^D2R^ calcium activity 3 seconds before (BE) and after (AE) the end of scratch (t-test, 1.320 ± 0.5270 *P = 0.0235, n = 17 trials).

Robust expression of GCaMP8s was found in the NAc LaSh^D1R^ neurons, as confirmed by amplified fluorescence achieved through immunostaining for GFP protein, with fiber placement just above the GCaMP8s-expressing neurons (Fig. 1B). The increase in Ca2+ activity in the NAc LaSh^D1R^ neurons during scratching is captured by changes in the GCaMP8s dynamics, which increases its fluorescence upon binding to Ca2+, allowing detection through the fiber photometry system (Fig. 1C). To probe the relationship between the NAc LaSh^D1R^ neurons’ calcium transient and chloroquine-induced scratching, we aligned the fluorescence signal with the start and end of scratching bouts. We found that GCaMP8s activity rises at the start of scratching (Fig. 1D-G), whereas no significant changes were detected at the end of scratching (Fig. 1I and J). We also observed an increase in the area under the curve (AUC) before and after the start of scratching (Fig. 1G). In contrast, the AUC at the end of scratching remains unchanged (Fig. 1K). Thus, the activity of NAc LaSh^D1R^ neurons correlates with the initiation of scratching bouts, suggesting a role of these neurons in facilitating pruritogen-induced scratching behavior (Fig. 1H).

Next, we looked at the population neural activity of NAc LaSh^D2R^ neurons in the NAc LaSh (Fig. 1L) with the help of the fiber photometry technique. We expressed the GCaMP8s sensor under the D2R promoter, using a similar strategy as with NAc LaSh^D1R^ neurons (Fig. 1A). In contrast to the NAc LaSh^D1R^ neurons, we found that the activity of NAc LaSh^D2R^ neurons do not correlate with the start of scratching bouts induced by choloroquine injected in the nape of the neck (Fig. 1M-P). Moreover, unlike the NAc LaSh^D1R^ neurons, the activity of the NAc LaSh^D2R^ rose prior to termination of the scratching bouts (Fig. 1R-T). Altogether, we found that the activity of the NAc LaSh^D1R^ ^and^ ^D2R^ neurons corresponds to distinct phases of itch-bouts induced by choloroquine.

Chronic itch caused by peripheral inflammation is associated with peripheral and central sensitization, leading to enhanced neuronal excitability to pruritic and non-pruritic stimuli (Ikoma et al., 2006; Li et al., 2023; Mahmoud et al., 2023; Guo et al., 2024). Hence, we next investigated the neural dynamics of NAc LaSh^D1R^ and NAc LaSh^D2R^ neurons during chronic itch using fiber photometry (Fig. 2A). We used the imiquimod-induced psoriasis model to induce chronic itch in mice (Sakai et al., 2016; Prajapati et al., 2026). Mice implanted with fiber-optic cannulas received daily topical imiquimod applications on the rostral back for six consecutive days, which induced psoriasis-like skin lesions and spontaneous scratching behavior (Fig. 2B). First, we recorded the spontaneous scratching behavior of mice, along with calcium transients of the NAc LaSh^D1R^ neurons. Our fiber photometry recordings revealed that the activity of NAc LaSh^D1R^ neurons rises at the start of scratching (Fig. 2C-F), resembling the response observed during chloroquine (CQ)-induced scratching (Fig. 1D-F). Under chronic itch conditions, NAc LaSh^D1R^ activity also showed a positive correlation with scratching duration (Fig. 2G), similar to what we observed in the acute itch model (Fig. 1G). Additionally, we found no change in calcium activity at the end of scratching (Fig. 2H-J), again consistent with observations from acute itch-induced scratching (Fig. 1H-J). Next, we examined NAc LaSh^D2R^ neuronal activity during chronic itch (Fig. 2K-R) and found that it resembles the response seen during CQ-induced scratching (Fig. 1K-S). Altogether, these results indicate that both NAc LaSh^D1R^ ^and^ ^D2R^ neurons respond to acute and chronic itch in a similar manner.

**Fig. 2.**
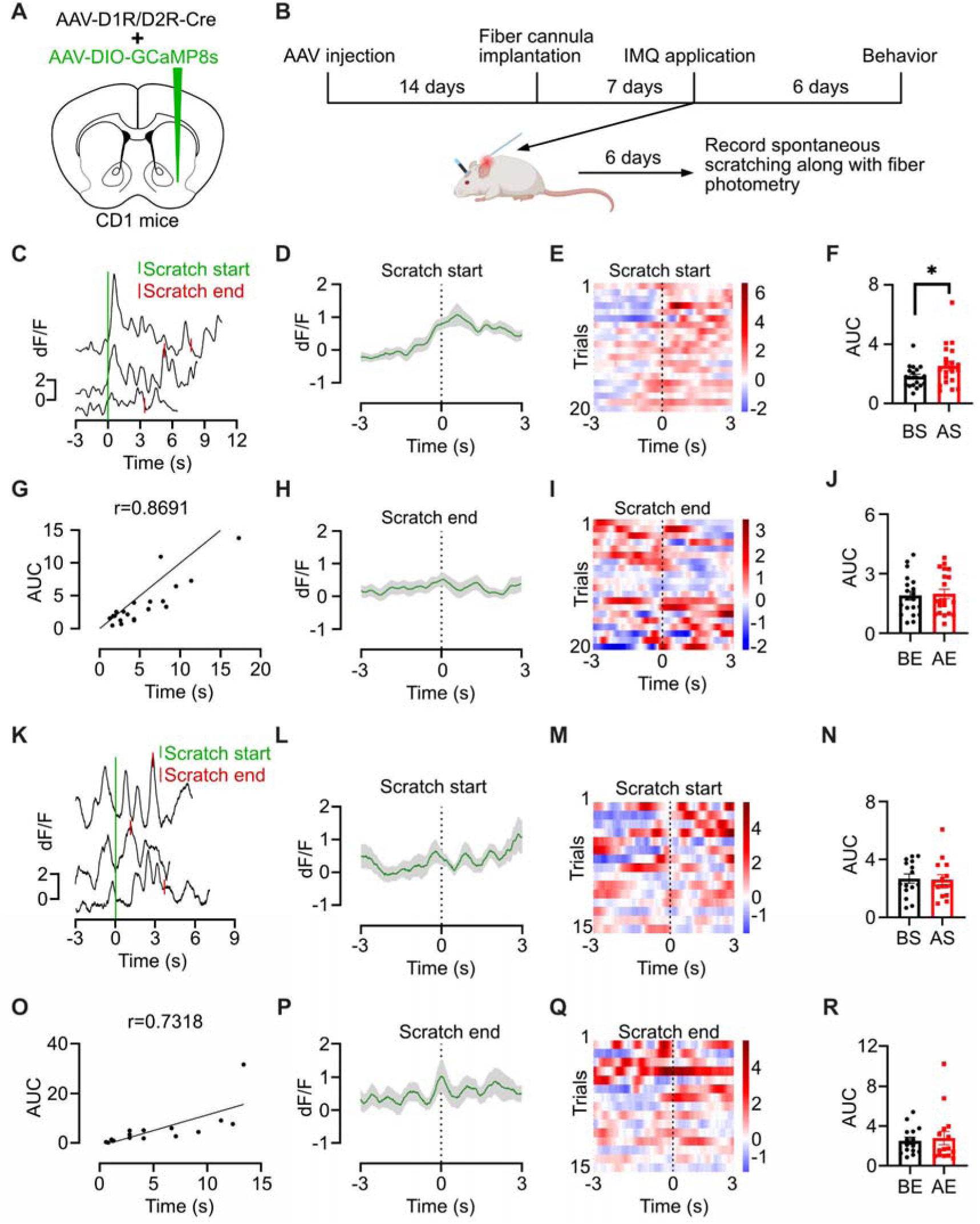
Engagement patterns of NAc LaSh^D1R^ and NAc LaSh^D2R^ neurons during chronic psoriatic itch. (A) Schematic of virus injection in the NAc LaSh. AAV-D1R-Cre or AAV-D2R-Cre, and AAV encoding Cre-dependent GCaMP8s were injected into the NAc LaSh of CD1 mice. (B) Schematic of the experimental timeline for viral vector expression and induction of chronic psoriatic itch using imiquimod. Subsequent spontaneous scratching behavior was recorded along with fiber photometry. (C) Representative ΔF/F traces of NAc LaSh^D1R^ neurons from three trials aligned to the start of chronic psoriatic itch-evoked spontaneous scratching. (D) The average fluorescence from NAc LaSh^D1R^ neurons at the start of scratch, with shaded area indicating SEM (n = 20 trials from 2 mice). (E) Heatmap of the fluorescence signal from NAc LaSh^D1R^ neurons at the start of scratch. (F) Quantification of area under the curve (AUC) of NAc LaSh^D1R^ activity 3 seconds before (BS) and after (AS) the start of scratch (t-test, 0.7309 ± 0.2709 *P = 0.0143, n = 20 trials). (G) Correlation of AUC of NAc LaSh^D1R^ activity with scratching duration under chronic psoriatic itch condition (****P < 0.0001). (H) The average fluorescence from NAc LaSh^D1R^ neurons at the end of scratch, with shaded area indicating SEM (n = 20 trials from 2 mice). (I) Heatmap of the fluorescence signal from NAc LaSh^D1R^ neurons at the end of scratch. (J) Quantification of area under the curve (AUC) of NAc LaSh^D1R^ activity 3 seconds before (BS) and after (AS) the end of scratch. (K) Representative ΔF/F traces of NAc LaSh^D2R^ neurons from three trials aligned to the start of chronic psoriatic itch-evoked spontaneous scratching. (L) The average fluorescence from NAc LaSh^D2R^ neurons at the start of scratch, with shaded area indicating SEM (n = 15 trials from 2 mice). (M) Heatmap of the fluorescence signal from NAc LaSh^D2R^ neurons at the start of scratch. (N) Quantification of area under the curve (AUC) of NAc LaSh^D2R^ activity 3 seconds before (BS) and after (AS) the start of scratch. (O) Correlation of AUC of NAc LaSh^D2R^ activity with scratching duration under chronic psoriatic itch condition (**P = 0.0013). (P) The average fluorescence from NAc LaSh^D2R^ neurons at the end of scratch, with the shaded area indicating SEM (n = 15 trials from 2 mice). (Q) Heatmap of the fluorescence signal from NAc LaSh^D2R^ neurons at the end of scratch. (R) Quantification of the area under the curve (AUC) of NAc LaSh^D2R^ activity 3 seconds before (BS) and after (AS) the end of scratch.

### Dopamine dynamics in NAc LaSh during acute and chronic itch-induced scratching behavior

Recent studies have demonstrated that VTA dopaminergic neurons are activated during scratching, with increased dopaminergic terminal activity in the lateral shell of the NAc compared to the medial shell (Yuan et al., 2018; Su et al., 2019). Despite these findings, the mechanisms underlying the emergence of dopamine activity during chronic itch remain unexplored. Hence, to delve into the dopamine dynamics during acute itch and how it emerges under chronic psoriatic itch-induced scratching, we utilized the dLight1.2 (Patriarchi et al., 2018). dLight1.2 is a GRAB (GPCR-activation-based) sensor, developed by coupling the human dopamine receptor 1 (DRD1) with the circularly permuted eGFP. When dopamine binds to the DRD1 receptor, it induces a conformational change in the circularly permuted eGFP, leading to an increase in fluorescent intensity that can be recorded using a fiber photometry system (Gunaydin et al., 2014; Kim et al., 2016; Patriarchi et al., 2018). We stereotaxically injected AAV-D1R-Cre and Cre-dependent dopamine sensor, DIO-dLight1.2, into the NAc LaSh of CD1 mice (Fig. 3A). We confirmed the expression of dLight1.2 and correct fiber placement after post-hoc staining for GFP (Fig. 3B). We found that the dopamine release sensed by NAc LaSh^D1R^ neurons rapidly increases at the onset of scratching, peaks at the end, and strongly correlates with the scratching duration (Fig. 3D-K). We observed an increase in AUC at the onset of scratching (Fig. 3G), but no change in AUC at the end of scratching bouts (Fig. 3K). This is because dopamine levels rise sharply and then fall rapidly at the bout’s termination, resulting in no net change in AUC at the end of scratching. The increase in dopamine levels at the start of scratching is consistent with a previous report (Liang et al., 2022). We further examined dopamine dynamics during chronic itch-induced scratching, as aberrant dopamine signaling may contribute to the persistent itch-scratch cycle observed in chronic itch conditions (Ishiuji, 2019; Lipman and Yosipovitch, 2021). Thus, next, we induced chronic itch in mice by repeated imiquimod applications and recorded their spontaneous scratching behavior, along with dopamine activity (Fig. 3L). After aligning the dopamine activity with the start and end of the scratching, we found that the pattern of dopamine release detected by NAc LaSh^D1R^ neurons during chronic itch was similar (increase in activity) to that seen during acute itch-induced scratching (Fig. 3M-S). Thus supporting our earlier finding that acute and chronic itch are processed similarly in the NAc LaSh.

**Fig. 3.**
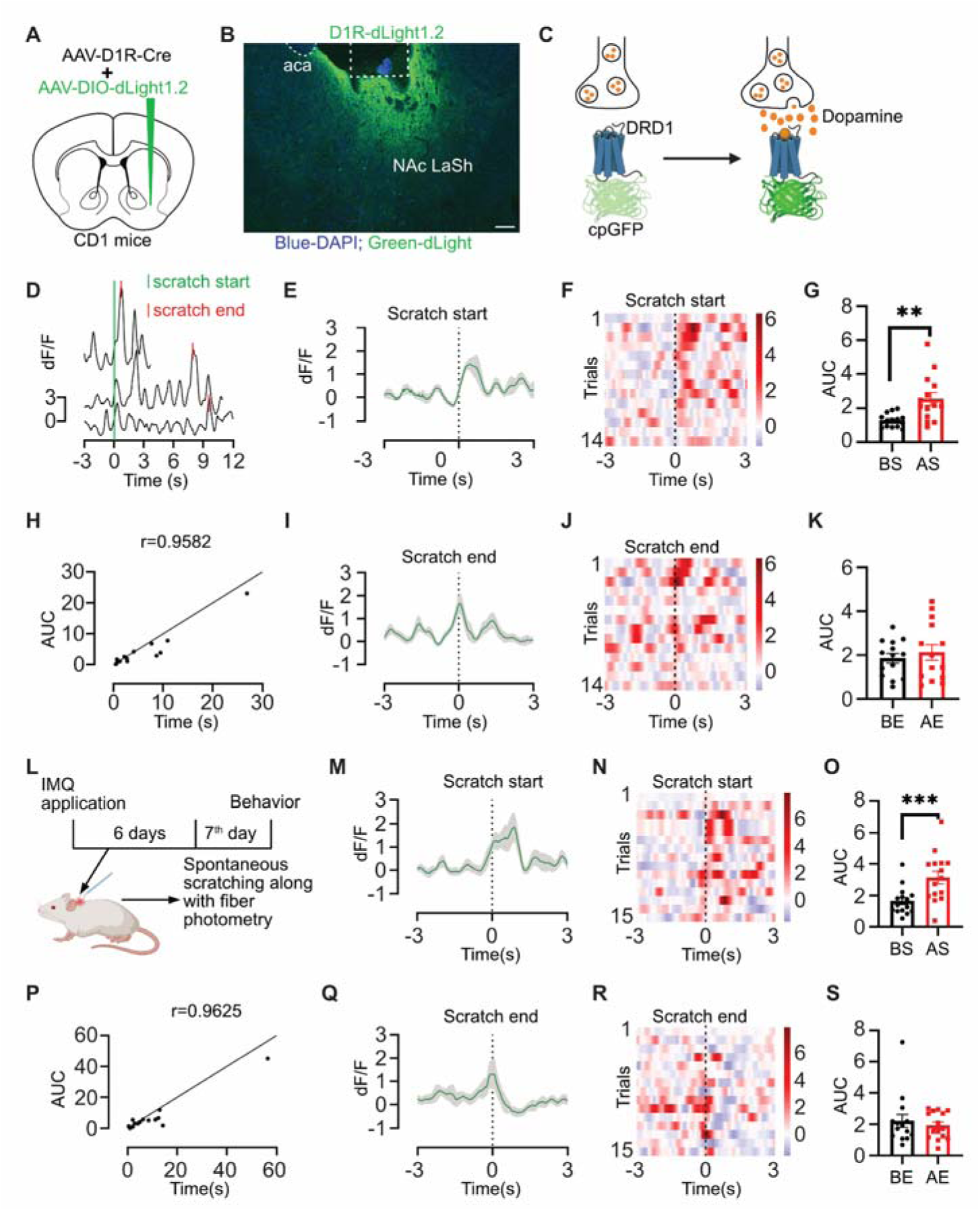
Dopamine dynamics detected by NAc LaSh^D1R^ neurons during acute and chronic psoriatic itch-evoked scratching. (A) Schematic of virus injection in the NAc LaSh. AAV-D1R-Cre and AAV encoding Cre-dependent dLight1.2 were injected into the NAc LaSh of CD1 mice. (B) Coronal section image of the NAc LaSh showing the D1R-dependent expression of AAV-DIO-dLight1.2 and placement of fiber optic cannula in NAc LaSh. Scale bar 100 µm. (C) Schematic showing the mechanism of action of the dLight1.2 sensor. (D) Representative ΔF/F traces of dopamine signal at NAc LaSh^D1R^ neurons from three trials aligned to the start of chloroquine-evoked scratching. (E) The average dopamine activity detected by NAc LaSh^D1R^ neurons at the start of chloroquine-evoked scratch, with shaded area indicating SEM (n = 14 trials from 2 mice). (F) Heatmap of the dopamine signal detected by NAc LaSh^D1R^ neurons at the start of chloroquine-evoked scratch. (G) Quantification of area under the curve (AUC) of dopamine activity 3 seconds before (BS) and after (AS) the start of chloroquine-evoked scratch (t-test, 1.255 ± 0.3641 **P = 0.0043, n = 14 trials). (H) Correlation of AUC of dopamine activity detected by NAc LaSh^D1R^ neurons with scratching duration (****P < 0.0001). (I) The average dopamine activity detected by NAc LaSh^D1R^ neurons at the end of the chloroquine-evoked scratch, with the shaded area indicating. (J) Heatmap of the dopamine signal detected by NAc LaSh^D1R^ neurons at the end of chloroquine-evoked scratch. (K) Quantification of the area under the curve (AUC) of dopamine activity 3 seconds before (BE) and after (AE) the end of the chloroquine-evoked scratch. (L) Schematic showing induction of chronic psoriatic itch and experimental paradigm. (M) The average dopamine activity detected by NAc LaSh^D1R^ neurons at the start of chronic psoriatic itch-evoked scratch, with shaded area indicating SEM (n = 15 from 2 mice). (N) Heatmap of the dopamine signal detected by NAc LaSh^D1R^ neurons at the start of chronic psoriatic itch-evoked scratch. (O) Quantification of area under the curve (AUC) of dopamine activity 3 seconds before (BS) and after (AS) the start of chronic psoriatic itch-evoked scratch t-test, 1.508 ± 0.3594 ***P = 0.0009, n = 15 trials). (P) Correlation of AUC of dopamine activity detected by NAc LaSh^D1R^ neurons with scratching duration under chronic psoriatic itch (****P < 0.0001). (Q) The average dopamine activity detected by NAc LaSh^D1R^ neurons at the end of chronic psoriatic itch-evoked scratch, with shaded area indicating SEM (n = 15 from 2 mice). (R) Heatmap of the dopamine signal detected by NAc LaSh^D1R^ neurons at the end of chronic psoriatic itch-evoked scratch. (S) Quantification of area under the curve (AUC) of dopamine activity 3 seconds before (BE) and after (AE) the end of chronic psoriatic itch-evoked scratch.

### Chronic itch increases basal dopamine levels in NAc LaSh

Recent studies have revealed similarities between the brain mechanisms involved in chronic itch and addiction (Ishiuji, 2019; Lipman and Yosipovitch, 2021). Similar to addiction, aberrant dopamine signaling may be involved in the reinforcement of the itch-scratch cycle under chronic itch conditions. However, our fiber photometry recordings demonstrate that dopamine exhibits similar dynamics during both acute and chronic itch-induced scratching (Fig. 3). Notably, this lack of difference in dopamine activity between acute and chronic itch-induced scratching may arise from the limitation of the fluorescent sensor-coupled fiber photometry technique, which only measures relative changes in dopamine levels (Grienberger and Konnerth, 2012; Simpson et al., 2024; Lodder et al., 2025). Hence, to directly measure absolute changes in dopamine concentration during the transition from acute to chronic itch, we used the microdialysis technique (Chefer et al., 2009). We implanted the microdialysis cannula above the NAc LaSh in male Wistar rats and induced psoriasis by applying imiquimod on the rostral-back skin (Fig. 4A and B). Next, the concentrations of dopamine (DA) and its metabolite (DOPAC) were assayed in dialysates collected from control and chronic itch (imiquimod-treated) groups of rats using a microdialysis sampling technique and quantified by HPLC-ECD. We found that DA and DOPAC levels in the NAc LaSh of imiquimod-treated rats increased significantly at the time points tested (Fig. 4C-E). Suggesting that the basal dopamine levels are higher in chronic itch conditions than in naive conditions. In addition, our fiber photometry recordings revealed that dopamine activity rises during scratching and peaks at its end (Fig. 3). This led us to hypothesize that a specific change in dopamine concentration may signal the termination of scratching. We further propose that, in chronic itch, because basal dopamine levels are already elevated (Fig. 4), the additional dopamine release triggered by scratching may not be sufficient to achieve the change in dopamine concentration needed to stop scratching, resulting in a persistent cycle of scratching. To test this hypothesis, we used levodopa (L-DOPA) to artificially increase dopamine levels in NAc and examined its effect on scratching behavior in both acute and chronic itch (Goldstein et al., 1982; Harun et al., 2016). We intraperitoneally injected either 5 mg/kg or 10 mg/kg of L-DOPA and tested its effect on chloroquine-induced scratching at 15 min and 2.5 h (Fig. S2A). We observed that L-DOPA effectively reduced chloroquine-induced acute scratching at both time points (Fig. SB, D, and E). Next, we examined the effect of L-DOPA on chronic itch-induced scratching. Imiquimod was applied daily on the rostral back skin of the mice for 6 consecutive days to induce chronic itch in mice. In contrast to acute itch, L-DOPA treatment did not affect the scratching in chronic itch (Fig. S2C and F). Therefore, dynamic changes in dopamine levels are required to terminate scratching.

**Fig. 4.**
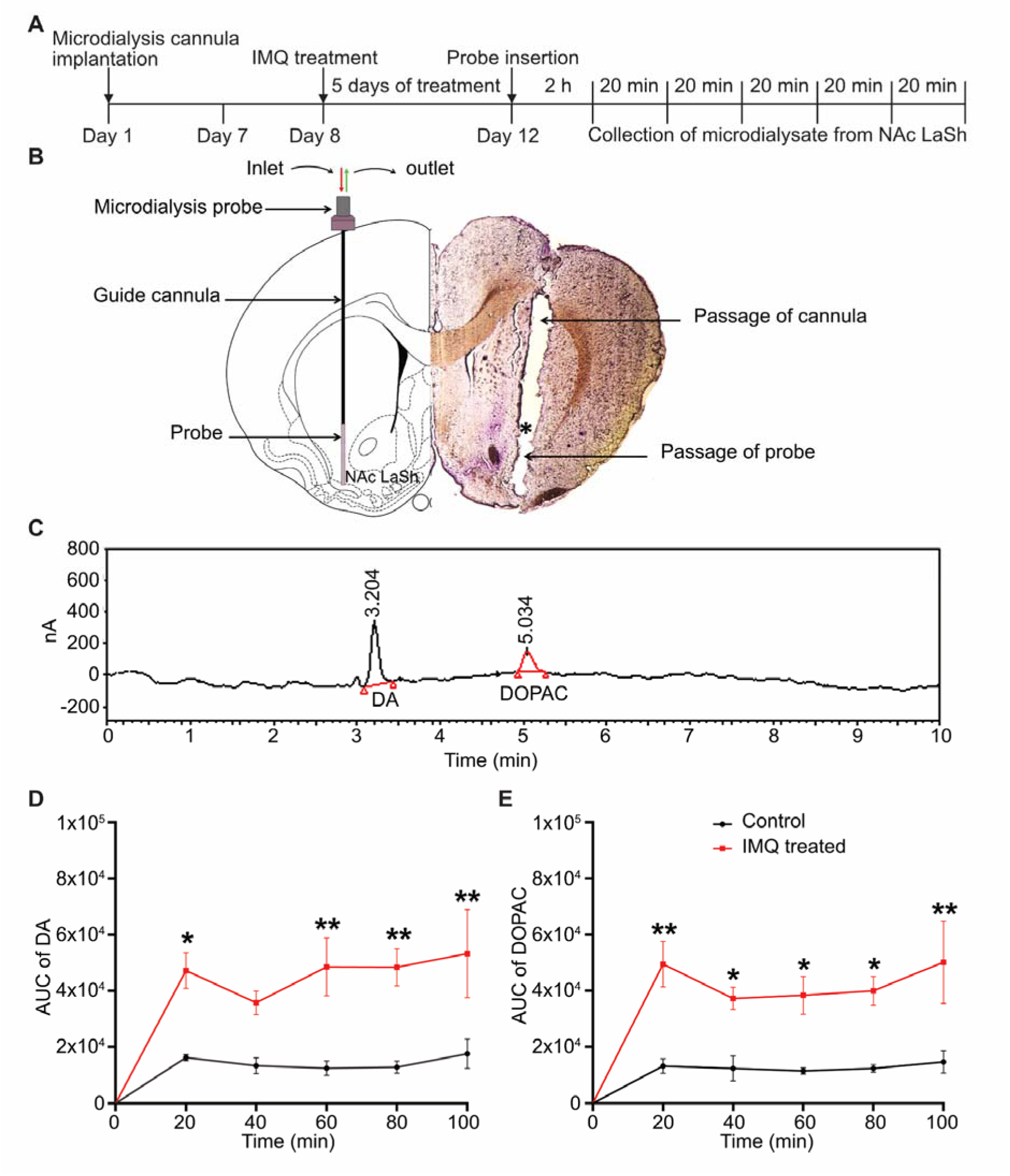
Chronic psoriatic itch basal DA and DOPAC levels in NAc LaSh in rats. (A) Schematic of the experimental timeline for microdialysis study. (B) Histological verification of microdialysis probe placement in NAc LaSh. (C) Chromatogram showed retention time (RT) of DA (RT: 3.204 min) and DOPAC (RT: 5.041) of the external standards. (D) Time course-AUC plot of DA measured every 20 minutes over a 100-minute sampling period in chronic psoriatic itch-induced rats (red) and the control (black) group (two-way ANOVA, p values from 20-100 minutes are 0.0163, 0.1503, 0.0037, 0.0043, and 0.0042, n=6 rats). (E) Time course-AUC plot of DOPAC measured every 20 minutes over a 100-minute sampling period in chronic psoriatic itch-induced rats (red) and the control (black) group (two-way ANOVA, p values from 20-100 minutes are 0.0011, 0.0436, 0.0240, 0.0191, and 0.0015, n=6 rats)

### NAc LaSh D1R and D2R neurons antagonistically modulate scratching behavior

Our fiber photometry revealed that NAc LaSh^D1R^ neurons increased their calcium transients at the start of scratching. In contrast, NAc LaSh^D2R^ neuronal activity increased at the end of scratching (Figs. 1 and 2), indicating opposite roles of NAc LaSh^D1R^ and NAc LaSh^D2R^ neurons in itch processing. Thus, next, we sought to determine how chemogenetic modulation of NAc LaSh^D1R^ and NAc LaSh^D2R^ neurons’ activity affects scratching under acute and chronic itch conditions. First, we co-injected AAV-D1R or D2R-Cre with AAV-DIO-hM3Dq into the NAc LaSh of CD1 mice (Fig. 5A) to chemogenetically activate neurons expressing either dopamine receptor. Robust hM3Dq expression was confirmed by mCherry labeling, which was activated readily upon DCZ administration, as confirmed by elevated cFos expression (Fig. 5B and G). As expected, activation of NAc LaSh^D1R^ neurons increased scratching (Fig. 5C-F), while activation of NAc LaSh^D2R^ neurons decreased scratching in both acute and chronic itch (Fig. 5H-K). Next, we employed the inhibitory DREADD, hM4Di, to suppress the activity of these same neuronal populations in the NAc LaSh (Fig. 6A, B, and G). We found that inhibition of NAc LaSh^D1R^ neurons suppressed scratching (Fig. 6C-F); however, inhibition of NAc LaSh^D2R^ neurons did not alter scratching behavior. Importantly, these effects are specific to the modulation of NAc LaSh^D1R^ and NAc LaSh^D2R^ neuronal activity, as DCZ administration did not affect scratching behavior in tdTomato-expressing control mice (Fig. S3). These chemogenetic results confirm that activation of NAc LaSh^D1R^ neurons promotes scratching, while activation of NAc LaSh^D2R^ neurons suppresses it.

**Fig. 5.**
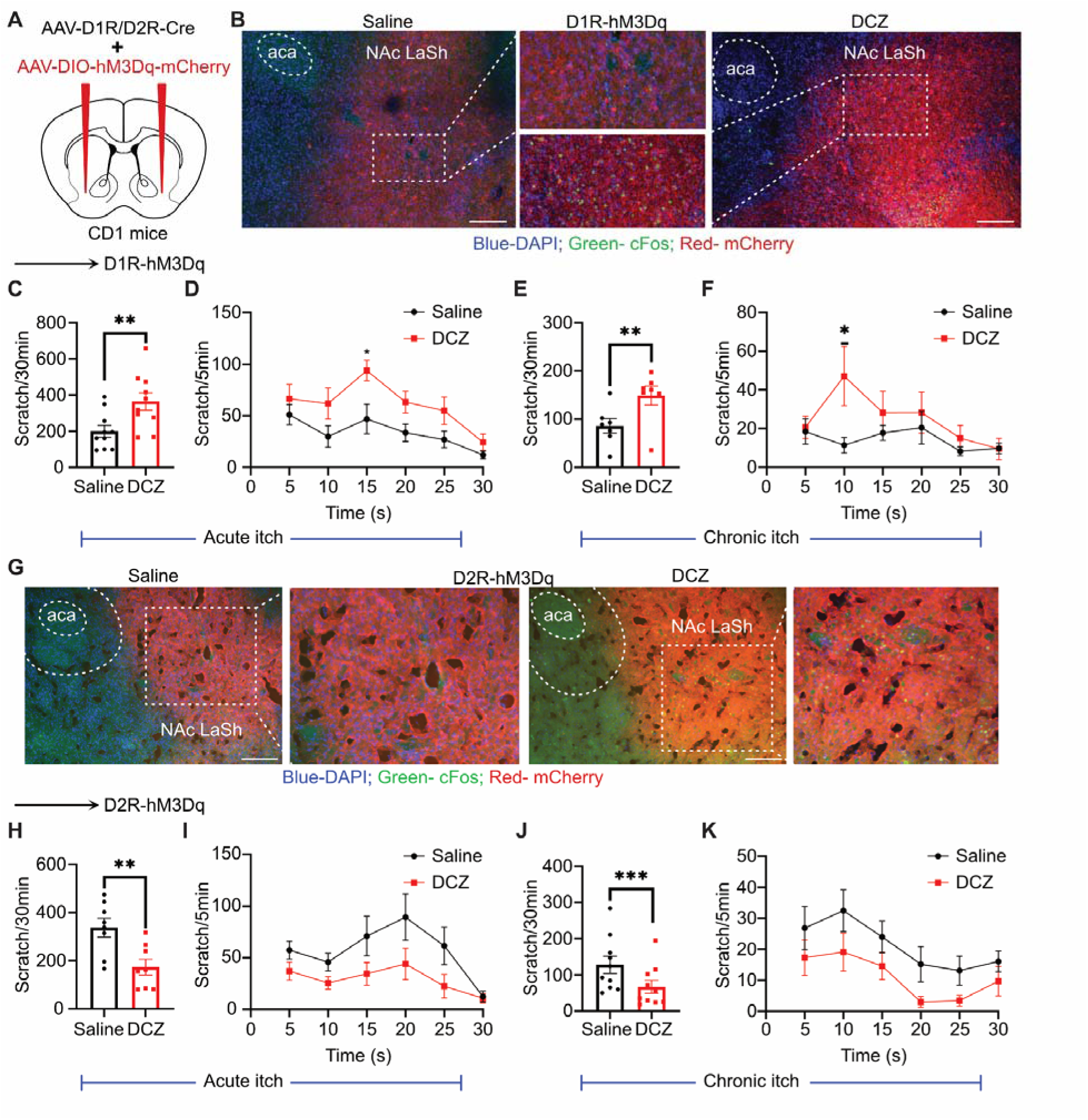
Chemogenetic activation of NAc LaSh^D1R^ and NAc LaSh^D2R^ neurons. (A) Schematic of the virus injection in NAc LaSh. AAV-D1R-Cre or AAV-D2R-Cre and AAV encoding Cre-dependent hM3Dq-mCherry were injected into the NAc LaSh of CD1 mice. (B) Coronal section image confirming the D1R-dependent expression of hM3Dq-mCherry in NAc LaSh and Fos (green) expression in hM3Dq-mCherry (red) expressing neurons and their overlap (yellow), post i.p.administration of DCZ compared with saline i.p.. Nuclei are stained with DAPI (blue). (C) Effect of DCZ/saline i.p. administration on chloroquine-induced scratching (t-test, 164.1± 44.95 **P = 0.0053, n=10 mice). (D) Temporal profile of scratching post DCZ/saline i.p. administration of chloroquine-induced scratching (two-way ANOVA, *P=0.0187). (E) Effect of DCZ/saline i.p. administration on chronic psoriatic itch-induced scratching (t-test, 62.86± 14.83 **P = 0.0054, n=7 mice). (F) Temporal profile of scratching post DCZ/saline i.p. administration of chronic psoriatic itch-induced scratching (two-way ANOVA, *P=0.0120). (G) Coronal section image confirming the D2R-dependent expression of hM3Dq-mCherry in NAc LaSh and Fos (green) expression in hM3Dq-mCherry (red) expressing neurons and their overlap (yellow), post i.p.administration of DCZ compared with saline i.p.. Nuclei are stained with DAPI (blue). (H) Effect of DCZ/Saline administration on chloroquine-induced scratching (t-test, 163.4± 41.33 **P = 0.0055, n=8 mice). (I) Temporal profile of scratching post DCZ/saline i.p. administration of chloroquine-induced scratching. (J) Effect of DCZ/saline i.p.administration on chronic psoriatic itch-induced scratching (t-test, 60.70± 12.57 ***P = 0.0009, n=10 mice). (K) Temporal profile of scratching post DCZ/saline i.p. administration of chronic psoriatic itch-induced scratching.

**Fig. 6.**
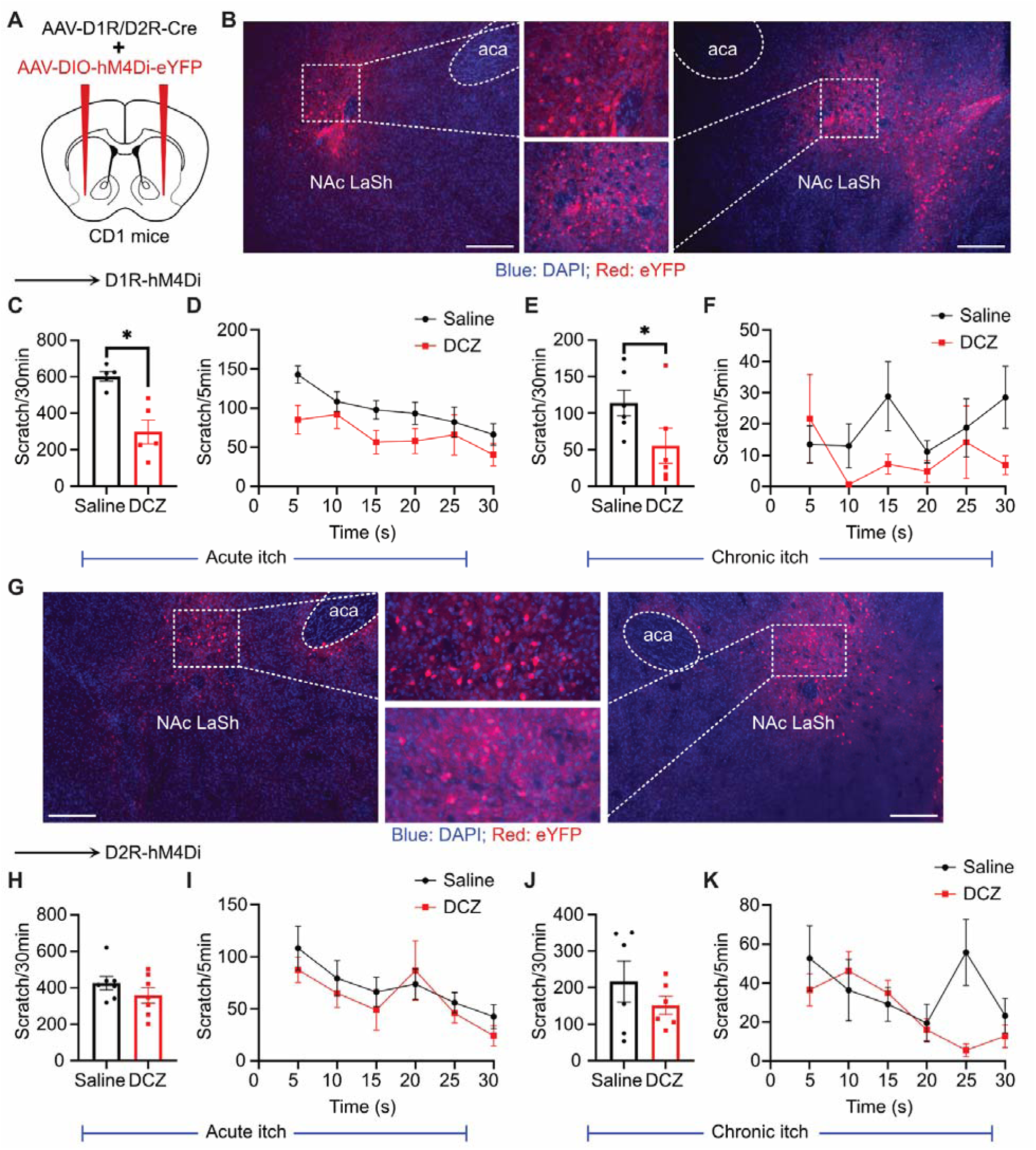
Chemogenetic inhibition of NAc LaSh^D1R^ and NAc LaSh^D2R^ neurons in NAc LaSh. (A) Schematic of the virus injection in NAc LaSh. AAV-D1R-Cre or AAV-D2R-Cre and AAV encoding Cre-dependent hM4Di-eYFP were injected into the NAc LaSh of CD1 mice. (B) Coronal section image confirming the D1R-dependent expression of hM4Di-eYFP (red blue-DAPI) in NAc LaSh. (C) Effect of DCZ/saline i.p. administration on chloroquine-induced scratching (t-test, 304.8± 71.22 *P = 0.0128, n=5 mice). (D) Temporal profile of scratching post DCZ/saline i.p. administration on chloroquine-induced scratching. (E) Effect of DCZ/saline i.p. administration on chronic psoriatic itch-induced scratching (t-test, 58.50± 20.28 *P = 0.0344, n=6 mice). (F) Temporal profile of scratching post DCZ/saline i.p. administration on chronic psoriatic itch-induced scratching. (G) Coronal section image confirming the D2R-dependent expression of hM4Di-eYFP (red blue-DAPI) in NAc LaSh. (H) Effect of DCZ/Saline administration on chloroquine-induced scratching. (I) Temporal profile of scratching post DCZ/saline i.p. administration on chloroquine-induced scratching. (J) Effect of DCZ/saline i.p. administration on chronic psoriatic itch-induced scratching. (K) Temporal profile of scratching post DCZ/saline i.p. administration on chronic psoriatic itch-induced scratching.

To further corroborate these findings, we employed optogenetics to activate NAc LaSh^D1R^ and NAc LaSh^D2R^ neurons. To that end, we injected the AAV-D1R or D2R-Cre viral vector, along with AAV-DIO-ChR2-mCherry, into the NAc LaSh of CD1 mice (Fig. 7A). Next, we implanted bilateral optical cannula just above the NAc LaSh and mice were allowed to recover for a week (Fig. 7C). During behavioral testing blue light (470 nm) was shone through the cannula in alternating 5-minute OFF and ON periods (Fig. 7B). As seen with chemogenetic activation (Fig. 6), light-driven stimulation of NAc LaSh^D1R^ neurons increased scratching during the ON periods compared to OFF (Fig. 7D and E), whereas control mice expressing eGFP showed no light-dependent changes (Fig. 7F and G). Conversely, optogenetic activation of NAc LaSh^D2R^ neurons decreased scratching (Fig. 7I and J), with no effect in eGFP controls (Fig. 7K and L). Altogether, we found that NAc LaSh^D1R^ and NAc LaSh^D2R^ neurons antagonistically drive scratching behavior in mice.

**Fig. 7.**
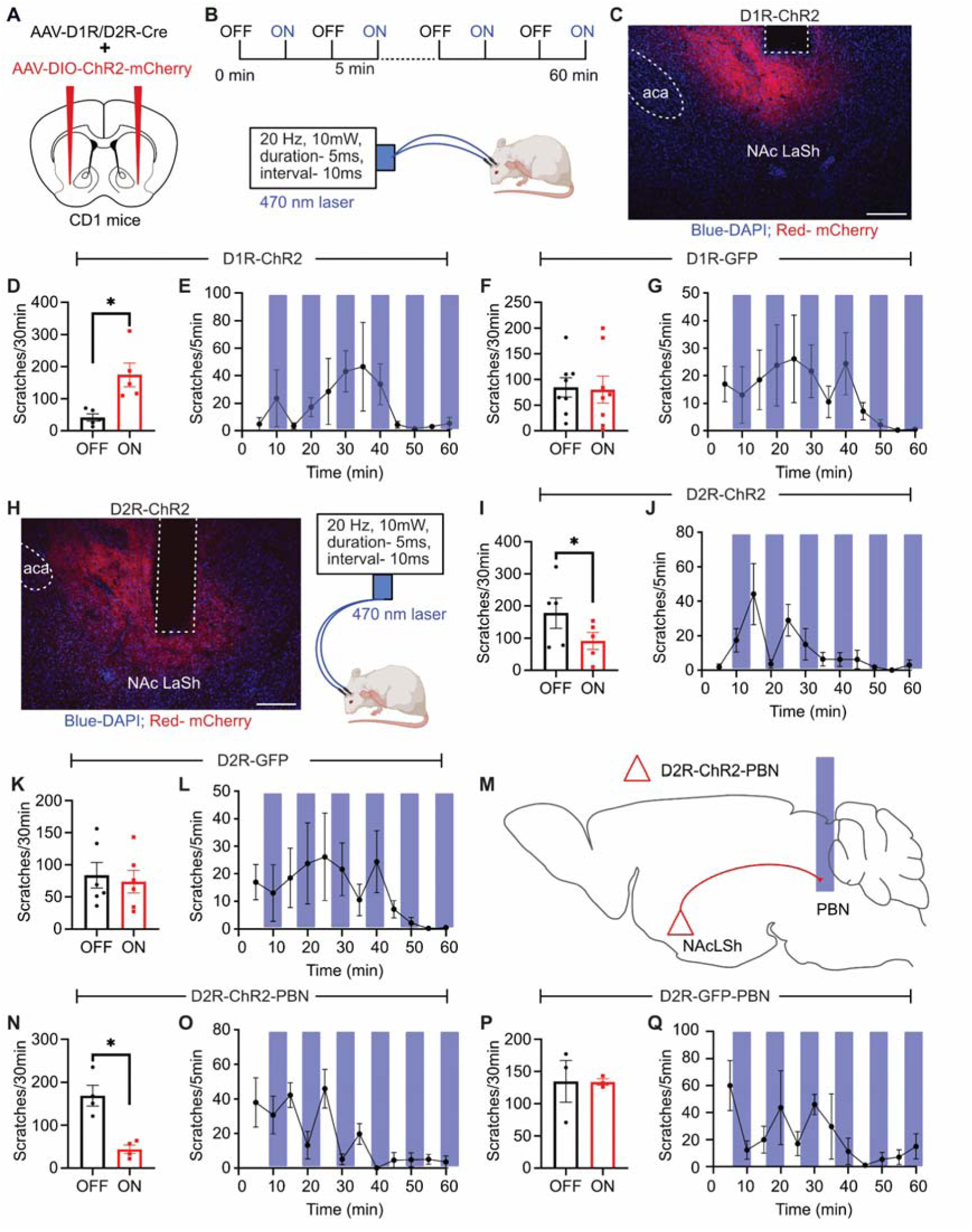
Optogenetic activation of NAc LaSh^D1R^ and NAc LaSh^D2R^ neurons in NAc LaSh. (A) Schematic of the virus injection in NAc LaSh. AAV-D1R-Cre or AAV-D2R-Cre and AAV encoding Cre-dependent ChR2-mCherry or Cre-dependent GFP were injected into the NAc LaSh of CD1 mice. (B) Schematic of the experimental paradigm. (C) Coronal section image of the NAc LaSh showing the D1R-dependent expression of ChR2-mCherry (red, blue-DAPI) in NAc LaSh. (D) Comparison of scratching behavior during light ON versus light OFF conditions post chloroquine administration in NAc LaSh^D1R-ChR2^-expressing mice (t-test, 132.4 ± 35.62 *P = 0.0205, n=5 mice). (E) Temporal profile of scratching across the alternating light ON/OFF epochs post chloroquine administration in NAc LaSh^D1R-ChR2^-expressing mice. (F) Comparison of scratching behavior during light ON versus light OFF conditions post chloroquine administration in NAc LaSh^D1R-GFP^-expressing mice. (G) Temporal profile of scratching across the alternating light ON/OFF epochs post-chloroquine administration in D1R-cre/GFP-expressing mice. (H) Coronal section image of the NAc LaSh showing the D2R-dependent expression of ChR2-mCherry (red, blue-DAPI) in NAc LaSh.. (I) Comparison of scratching behavior during light ON versus light OFF conditions post chloroquine administration in NAc LaSh^D2R-ChR2^-expressing mice (t-test, 43.0 ± 14.42 *P = 0.0406, n=5 mice). (J) Temporal profile of scratching across the alternating light ON/OFF epochs post chloroquine administration in NAc LaSh^D2R-ChR2^-expressing mice. (K) Comparison of scratching behavior during light ON versus light OFF conditions post chloroquine administration in NAc LaSh^D2R-GFP^-expressing mice. (L) Temporal profile of scratching across the alternating light ON/OFF epochs post-chloroquine administration in NAc LaSh^D2R-GFP^-expressing mice. (M) Schematic of the virus injection. AAV-D2R-Cre and AAV encoding Cre-dependent ChR2-mCherry or Cre-dependent GFP were injected into the NAc LaSh of CD1 mice. Optic fiber was implanted at PBN for terminal activation. (N) Comparison of scratching behavior during light ON versus light OFF conditions post chloroquine administration in NAc LaSh^D2R-ChR2-PBN^ mice (t-test, 124.8 ± 25.83 *P = 0.0169, n=4 mice). (O) Temporal profile of scratching across the alternating light ON/OFF epochs post chloroquine administration in NAc LaSh^D2R-ChR2-PBN^ mice. (P) Comparison of scratching behavior during light ON versus light OFF conditions post chloroquine administration in NAc LaSh^D2R-GFP-PBN^ mice. (Q) Temporal profile of scratching across the alternating light ON/OFF epochs post-chloroquine administration in NAc LaSh^D2R-GFP-PBN^ mice.

### NAc LaSh D2R neurons suppress scratching via their PBN terminals

We sought to determine the brain-wide projections of NAc LaSh^D1R^ and NAc LaSh^D2R^ neurons. We expressed synaptophysin-tagged GFP in NAc LaSh^D1R^ neurons by co-injecting AAV-D1R-Cre and AAV-DIO-mSynGFP (Fig. S4A). Synaptophysin is specifically enriched in synaptic vesicles of axon terminals; hence, the GFP-tagged version of the protein can be used for brain-wide projection-mapping of cell types of choice. We observed green puncta in several forebrain regions, including the frontal association cortex (FrA), secondary motor cortex, caudate putamen (CPu), and ventral pallidum (Fig. S4C, D, and E). In the midbrain, it projects to the lateral part of the VTA and the substantia nigra (SNR) (Fig. S4D and E). Next, we used the same approach to map the brain-wide projections of NAc LaSh^D2R^ neurons (Fig. S5). In the forebrain, NAc LaSh^D2R^ neurons project to the globus pallidus, ventral pallidum, entopeduncular nucleus, and lateral hypothalamus (Fig. S5C, D, and E). As seen in projection-mapping studies of NAc LaSh^D1R^ and NAc LaSh^D2R^ neurons, which project to the VTA and SNR (Yang et al., 2018). Interestingly, NAc LaSh^D2R^ and not NAc LaSh^D1R^ neurons project to the lateral parabrachial nucleus (LPBN) (Fig. S5D, bottom right). The LPBN acts as a critical hub in processing and transmitting itch information to various parts of the brain (Mu et al., 2017; Piyush Shah and Barik, 2022; Prajapati et al., 2024; Sun, 2025). Given the GABAergic nature of NAc LaSh^D2R^ neurons, we hypothesized that activating the LPBN terminals of the NAc LaSh^D2R^ neurons would inhibit PBN glutamatergic neurons to impair the itch transmission. To test this idea, we expressed ChR2 in NAc LaSh^D2R^ neurons, and a bilateral cannula was implanted just above the LPBN (Fig. 7M). We observed that stimulation of the LPBN terminals of NAc LaSh^D2R^ neurons decreased scratching (Fig. 7N and O), whereas stimulation of eGFP terminals did not affect scratching (Fig. 7P and Q). Therefore, we conclude that NAc LaSh^D2R^ neurons suppress scratching by inhibiting LPBN neurons.

## Discussion

Itch-induced scratching engages dopaminergic receptor-expressing neurons in the striatal NAc, yet how these neurons regulate the onset and offset of scratching bouts in acute and pathological itch remains unclear. Here, we found that NAc LaSh^D1R^ and NAc LaSh^D2R^ neurons exhibit distinct temporal profiles during scratching: NAc LaSh^D1R^ neurons showed increased calcium activity at the start of scratching, whereas NAc LaSh^D2R^ neurons became more active close to the end of the scratching bout (Figs. 1 and 2). Consistent with these opposing activity patterns, artificially manipulating these neurons had opposite effects on scratching behavior (Figs. 5,6, and 7). Interestingly, we found that the NAc LaSh^D1R^ and NAc LaSh^D2R^ neurons process acute and chronic itch in a similar fashion (Figs. 1 and 2). In parallel, we measured dopamine release in the NAc LaSh and found that it increased at the onset of scratching and peaked at its end (Fig. 3). Based on this, we hypothesized that a specific change in dopamine concentration is necessary to signal scratch termination. Notably, during chronic itch, overall dopamine levels in the NAc were elevated (Fig. 4), which may contribute to the persistent itch-scratch cycle by disrupting dopamine dynamics.

Using multiplexed in situ hybridization, we first showed that our transsectional viral genetic strategy efficiently labeled the D1R and D2R populations in NAc LaSh (Fig. S1), a finding that was crucial for our subsequent experiments. Notably, the efficiency of labeling D1R and D2R neurons in our experiments was comparable to that reported in recent studies (Liu et al., 2025b; Zhang et al., 2025). Using these viral vectors in combination with fiber photometry, we showed that NAc LaSh^D1R^ neuron activity increases at the onset of scratching and positively correlates with scratching duration (Fig. 1D-G), suggesting that NAc LaSh^D1R^ neuron activity is required to promote scratching behavior and may encode the motivational drive to scratch.

Moreover, chemogenetic and optogenetic stimulation data further support these findings (Figs. 5-7). Our results are consistent with previous findings showing that pharmacological inhibition of the D1R receptor and activation of D2R receptor reduces itch in mice (Akimoto and Furuse, 2011; Liang et al., 2022) and rats (Merali and Piggins, 1990).

Chronic pain is associated with a hypodopaminergic state (Jääskeläinen et al., 2001; Jarcho et al., 2012; Borsook et al., 2016; Yang et al., 2021). Itch and pain are known to be in an antagonistic relationship (Ikoma et al., 2006; Davidson and Giesler, 2010); thus, it is likely that chronic itch leads to increased dopamine concentration in the nucleus accumbens. A recent study has shown that a complex circuitry between the spinal cord dorsal, PBN, substantia nigra pars reticulata (SNR), and VTA is involved in the hypodopaminergic state under chronic pain conditions (Yang et al., 2021); however, the circuit mechanism for the hyperdopaminergic state in chronic itch conditions is not yet known. Mechanistically, this increase in dopamine concentration under chronic itch can be attributed to heightened, sustained firing of VTA dopaminergic neurons. Interestingly, recent studies in rodents have shown that chronic itch increases neuronal excitability across multiple brain regions (Li et al., 2023; Guo et al., 2024). However, the excitability of VTA dopaminergic neurons during chronic itch has not yet been tested. VTA dopaminergic neurons receive inputs from diverse brain regions, including PBN and LHA (Watabe-Uchida et al., 2012; Beier et al., 2015; Morales and Margolis, 2017). Firstly, PBN transmits itch-related signals to forebrain regions, along with the VTA (Mu et al., 2017; Chiang et al., 2019; Ren et al., 2023; Prajapati et al., 2024; Sun, 2025). Hence, it is possible that chronic itch sensitizes itch-processing glutamatergic neurons in the PBN, thereby increasing the activity of VTA dopaminergic neurons and elevating dopamine release in the nucleus NAc LaSh. The second potential mechanism could involve input from the lateral hypothalamic area (LHA) to the VTA. In a recent study, we found that psoriasis increases the firing of LHA stress-sensitive neurons that project to the VTA (Prajapati et al., 2026). This suggests that LHA stress-sensitive neurons may drive increased firing of VTA dopaminergic neurons, leading to elevated dopamine levels in the NAc LaSh. In the future, it will be interesting to examine which of these inputs to the VTA is important for driving the hyperdopaminergic state under chronic itch.

---

**Fig. S1.**
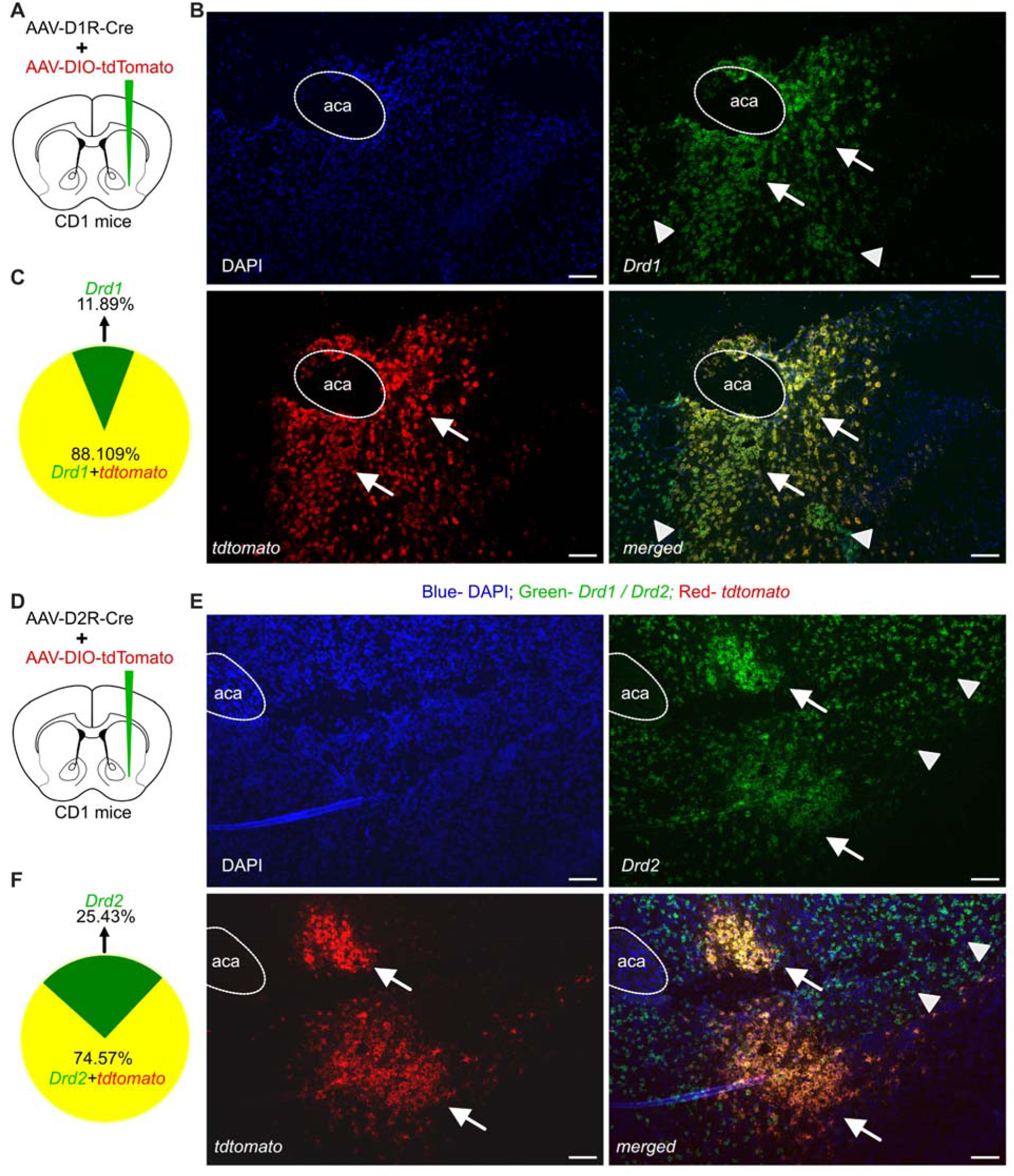
Efficiency of D1R-cre and D2R-cre viral vectors. (A) Schematic of the virus injection in NAc LaSh. AAV-D1R-Cre and AAV encoding Cre-dependent tdTomato were injected into the NAc LaSh of CD1 mice. (B) Validation of D1R-cre virus efficiency by Insitu hybridization. The top right panel shows Drd1 (green) mRNA expression, the bottom left panel shows tdTomato (red) expression, and the bottom right panel shows the overlap (yellow) of both. Nuclei are stained with DAPI (blue). (C) Pie chart showing the neuronal labeling distribution, yellow represents co-localization of Drd1 mRNA and tdTomato, and green represents the neurons labeled with Drd1 mRNA alone. (D) Schematic of the virus injection in NAc LaSh. AAV-D2R-Cre and AAV encoding Cre-dependent tdTomato were injected into the NAc LaSh of CD1 mice. (E) Validation of D2R-cre virus efficiency by Insitu hybridization. The top right panel shows Drd2 (green) mRNA expression, the bottom left panel shows tdTomato (red) expression, and the bottom right panel shows the overlap (yellow) of both. Nuclei are stained with DAPI (blue). (F) Pie chart showing the neuronal labeling distribution, yellow represents co-localization of Drd2 mRNA and tdTomato, and green represents the neurons labeled with Drd2 mRNA alone.

**Fig. S2.**
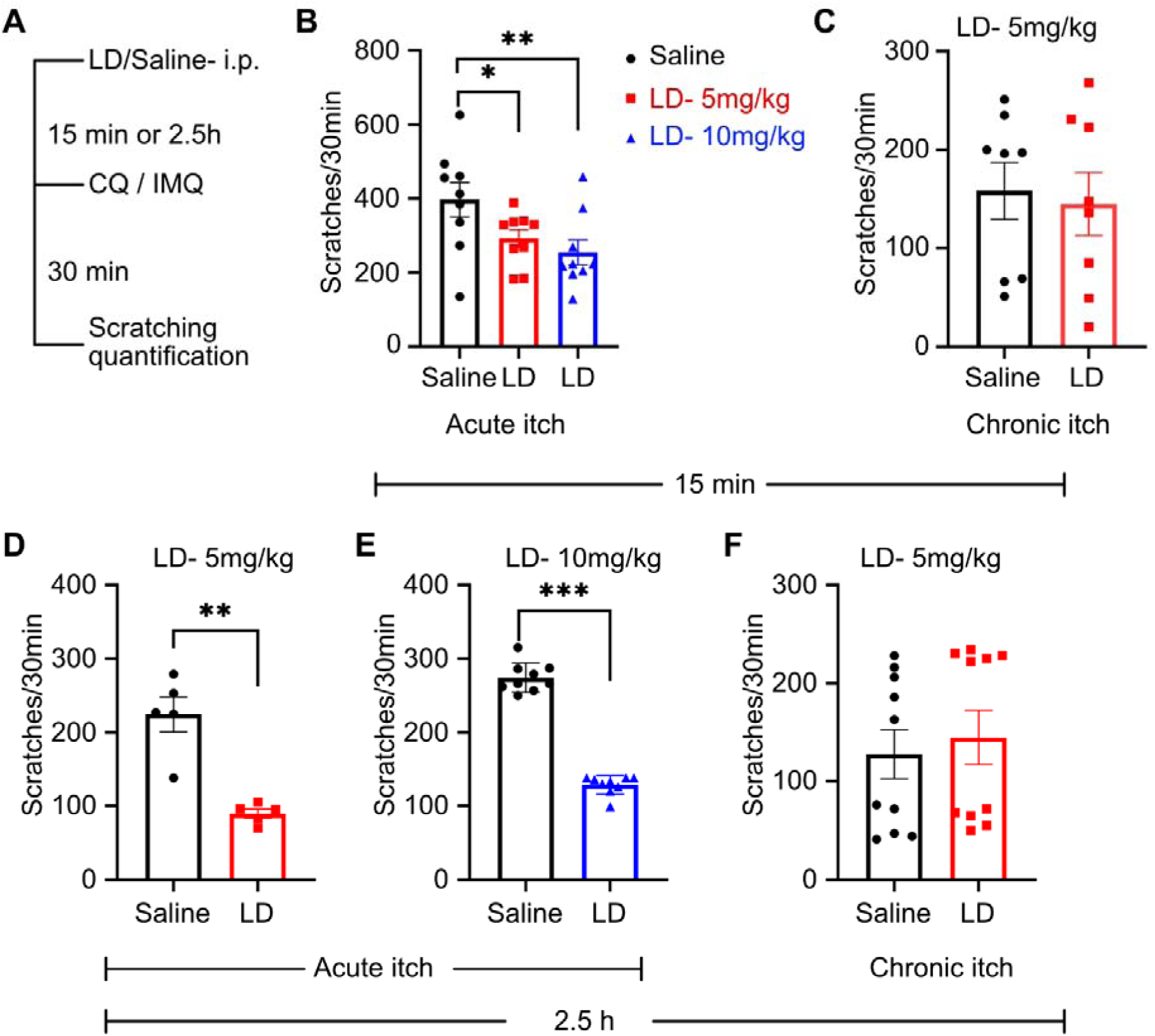
Effect of L-DOPA administration on acute and chronic psoriatic itch-induced scratching. (A) Schematic of the experimental paradigm (B) Effect of L-DOPA i.p. administration at dosage 5 or 10mg/kg body weight, 15 minutes prior to chloroquine injection, on scratching, compared with saline i.p. (two-way ANOVA, saline vs. 5 mg/Kg *P= 0.0449, saline vs. 10 mg/Kg **P=0.0071). (C) Effect of L-DOPA i.p. administration of dosage 5mg/kg, 15 minutes prior to behavioral testing, on chronic psoriatic itch-induced scratching compared with saline i.p. (D) Effect of L-DOPA i.p. administration of 5mg/kg, 2.5 hours prior to chloroquine injection, on scratching, compared with saline i.p (t-test,134.6 ± 23.70 **P = 0.0047, n=5 mice). (E) Effect of L-DOPA i.p. administration of 10mg/kg dosage 2.5 hours prior to chloroquine injection, on scratching, compared with saline i.p (t-test,145.6 ± 8.821 ****P<0.0001, n=9 mice). (F) Effect of L-DOPA i.p. administration of dosage 5mg/kg, 2.5 hours prior to behavioral testing, on chronic psoriatic itch-induced scratching compared with saline i.p.

**Fig. S3.**
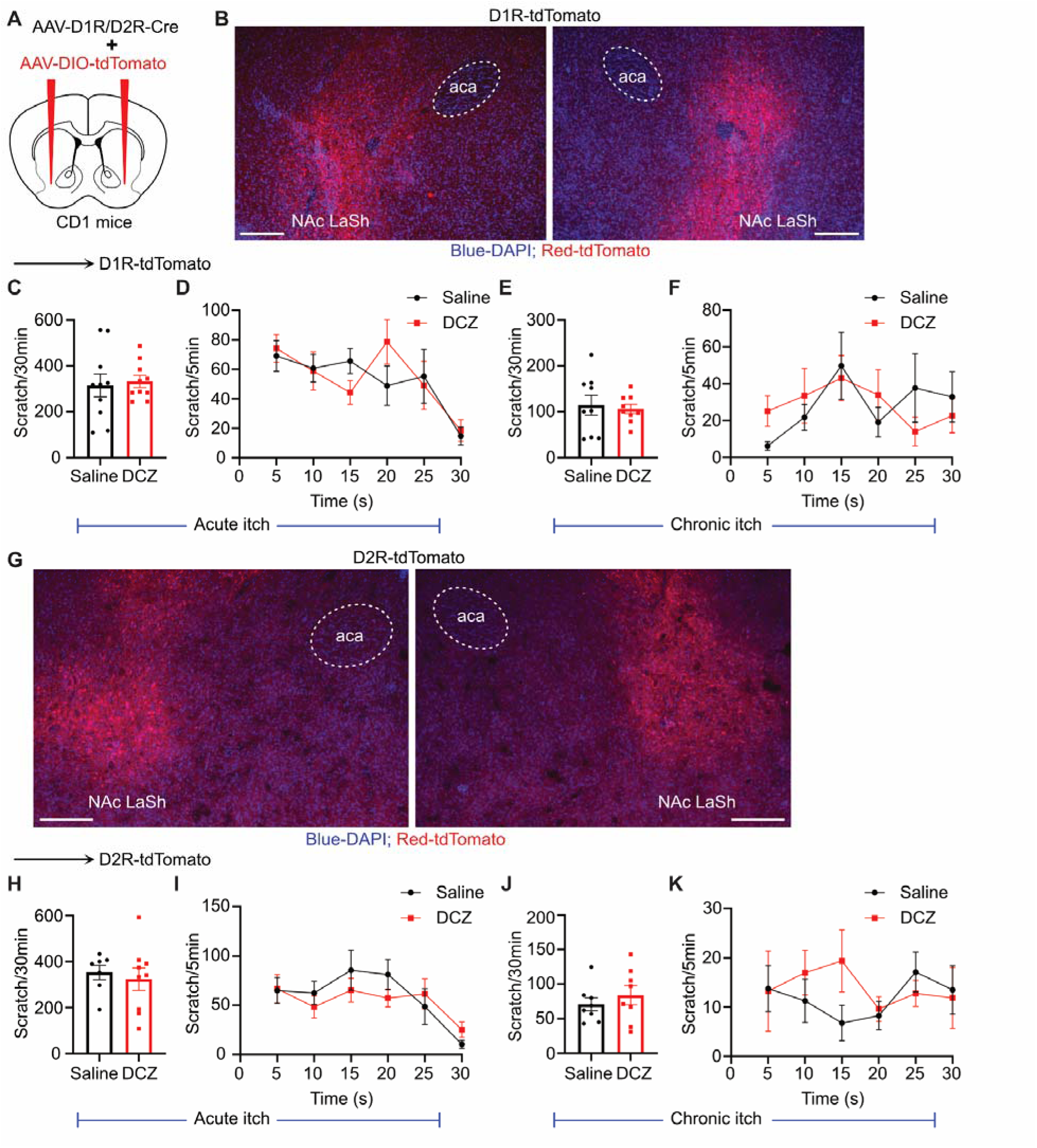
tdTomato control for chemogenetic manipulation of NAc LaSh^D1R^ and NAc LaSh^D2R^ neurons. (A) Schematic of the virus injection in NAc LaSh. AAV-D1R-Cre or AAV-D2R-Cre, with AAV encoding Cre-dependent tdTomato, was injected into the NAc LaSh of CD1 mice. (B) Coronal section image confirming the D1R-dependent expression of tdTomato (red blue-DAPI) in NAc LaSh. (C) Comparison of DCZ/saline i.p.administration on chloroquine-induced scratching in D1R-tdTomato expressing mice. (D) Temporal profile of scratching post DCZ/saline i.p. administration on chloroquine-induced scratching in D1R-tdTomato expressing mice. (E) Comparison of DCZ/saline i.p.administration on chronic psoriatic itch-induced scratching in D1R-tdTomato expressing mice. (F) Temporal profile of scratching post DCZ/saline i.p. administration of chronic psoriatic itch-induced scratching in D1R-tdTomato expressing mice. (G) Coronal section image confirming the D2R-dependent expression of tdTomato (red blue-DAPI) in NAc LaSh. (H) Comparison of DCZ/saline i.p.administration on chloroquine-induced scratching in D2R-tdTomato expressing mice. (I) Temporal profile of scratching post DCZ/saline i.p. administration of chloroquine-induced scratching in D2R-tdTomato expressing mice. (J) Comparison of DCZ/saline i.p.administration on chronic psoriatic itch-induced scratching in D2R-tdTomato expressing mice. (K) Temporal profile of scratching post DCZ/saline i.p. administration of chronic psoriatic itch-induced scratching in D2R-tdTomato expressing mice.

**Fig. S4.**
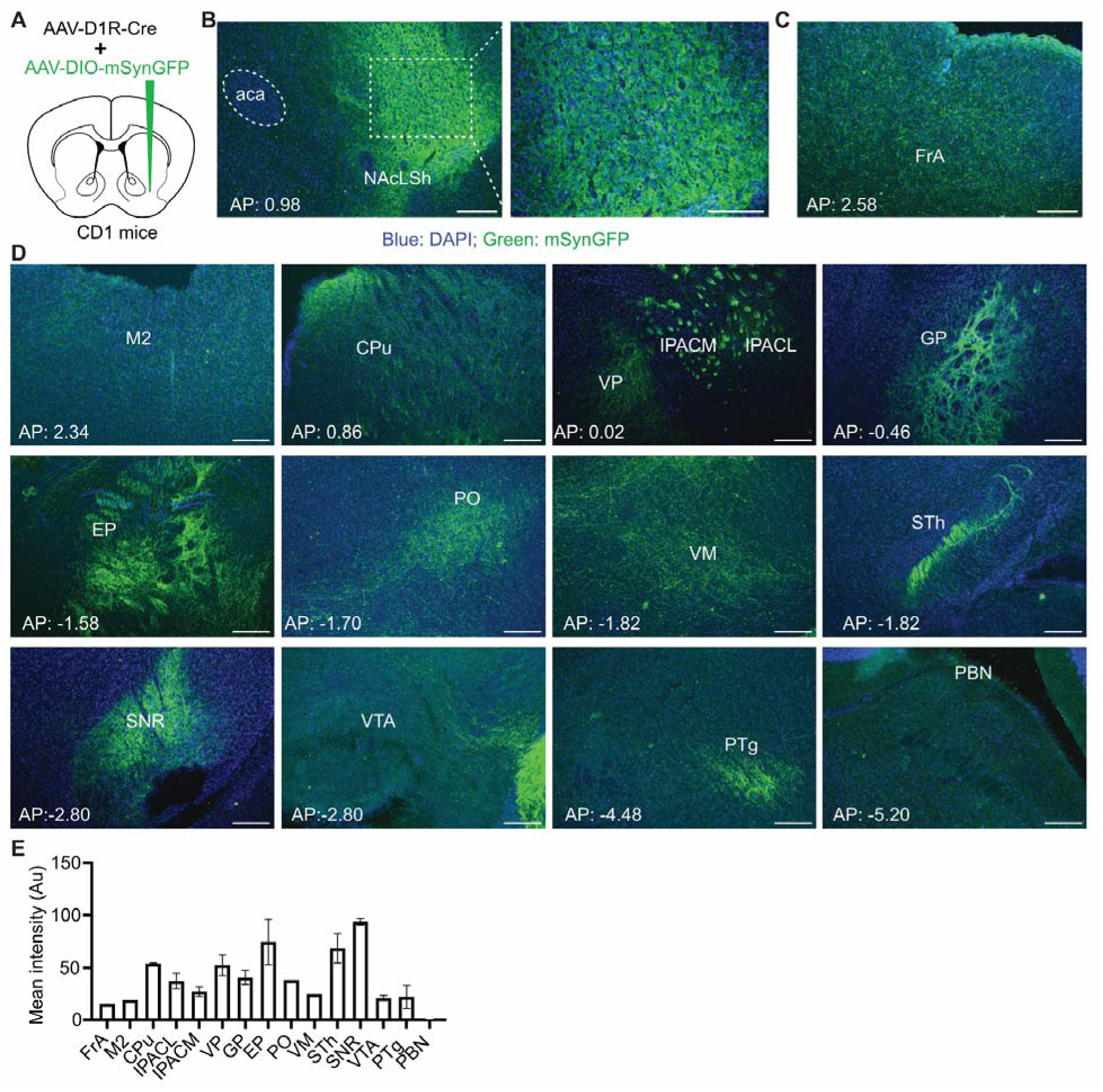
Projections of NAc LaSh^D1R^ neurons. (A) Schematic of the virus injection in NAc LaSh. AAV-D1R-Cre and AAV encoding Cre-dependent mSynGFP were injected into the NAc LaSh of CD1 mice. (B) Coronal section image confirming the D1R-dependent expression of tdTomato (red blue-DAPI) in NAc LaSh. (C-D) Brain regions receiving projections from NAc LaSh^D1R^ neurons. (E) Quantification of the mean intensity of projections from NAc LaSh^D1R^ neurons to various brain regions.

**Fig. S5.**
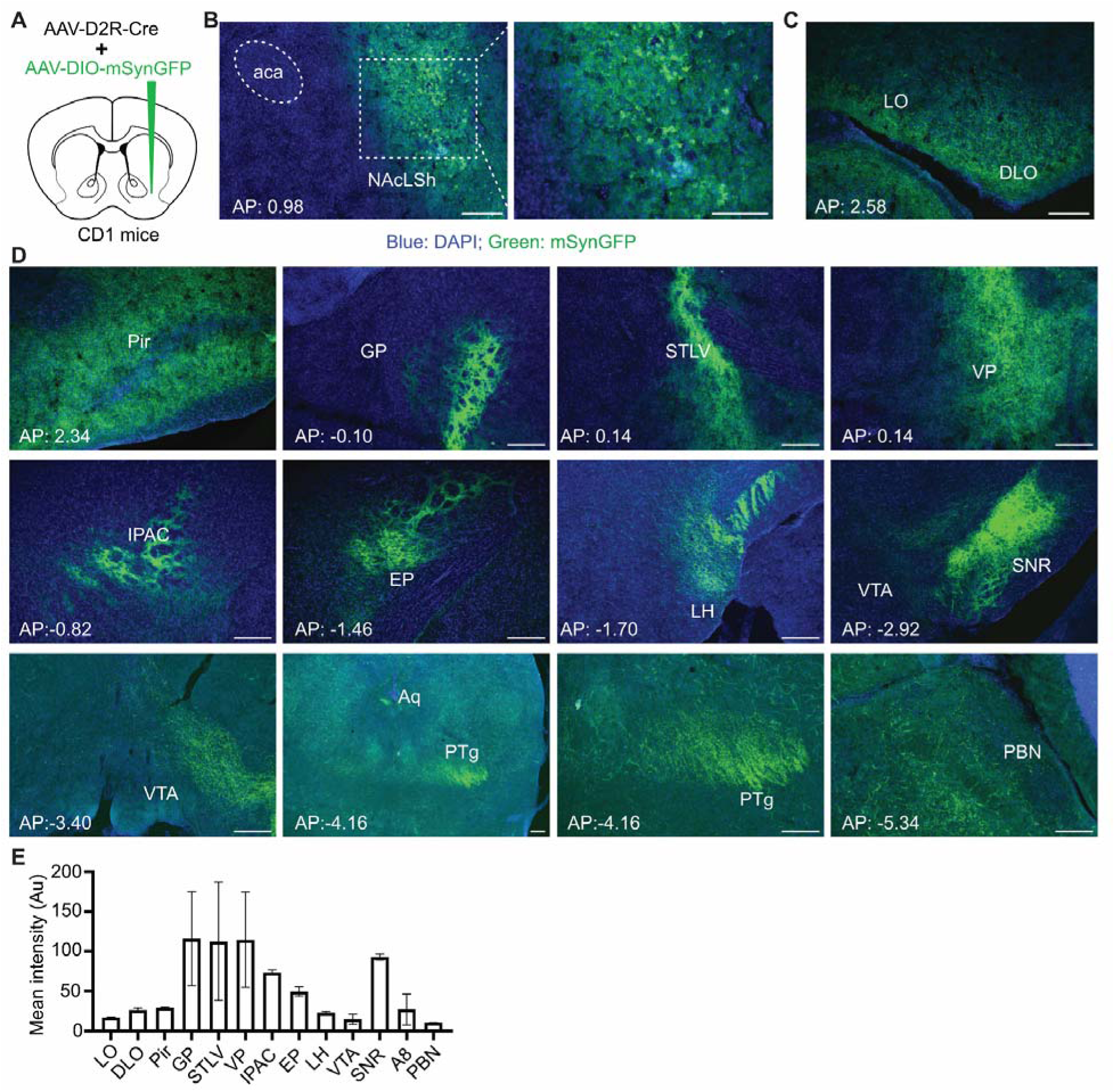
Projections of NAc LaSh^D2R^ neurons. (A) Schematic of the virus injection in NAc LaSh. AAV-D2R-Cre and AAV encoding Cre-dependent mSynGFP were injected into the NAc LaSh of CD1 mice. (B) Coronal section image confirming the D2R-dependent expression of tdTomato (red blue-DAPI) in NAc LaSh. (C-D) Brain regions receiving projections from NAc LaSh^D2R^ neurons. (E) Quantification of the mean intensity of projections from NAc LaSh^D2R^ neurons to various brain regions.

## Notes

### Competing Interest Statement

The authors have declared no competing interest.

### Funding

- Wellcome Trust/DBT India Alliance

